# Interrogation of SLFN11 in pediatric sarcomas uncovers an unexpected biological role and a novel therapeutic approach to overcoming resistance to replicative stress

**DOI:** 10.1101/2020.11.05.370155

**Authors:** Jessica Gartrell, Marcia Mellado-Largarde, Nancy E. Martinez, Michael R. Clay, Armita Bahrami, Natasha Sahr, April Sykes, Kaley Blankenship, Lauren Hoffmann, Jia Xie, Hyekyung Plumley, Nathaniel Twarog, Michele Connelly, Koon-Kiu Yan, Jiyang Yu, Shaina N. Porter, Shondra M. Pruett-Miller, Geoffrey Neale, Christopher L. Tinkle, Sara M. Federico, Elizabeth A. Stewart, Anang A. Shelat

## Abstract

Pediatric sarcomas represent a heterogeneous group of malignancies that exhibit variable response to DNA damaging chemotherapy. Schlafen family member 11 protein (SLFN11) increases sensitivity to replicative stress, and *SLFN11* gene silencing has been implicated as a common mechanism of drug resistance in tumors in adults. We found SLFN11 to be widely expressed in our cohort of pediatric sarcomas. In sarcoma cell lines, protein expression strongly correlated with response to the PARP inhibitor talazoparib (TAL) and the topoisomerase I inhibitor irinotecan (IRN), with SLFN11 knockout resulting in significant loss of sensitivity *in vitro* and *in vivo.* However, SLFN11 expression was not associated with favorable outcomes in a retrospective analysis of our patient cohort; instead, the protein was retained and promoted tumor growth and evasion. Furthermore, we show that pediatric sarcomas develop resistance to TAL and IRN through impaired intrinsic apoptosis, and that resistance can be reversed by selective inhibition of BCL-XL.

**Statement of Significance:** The role of SLFN11 in pediatric sarcomas has not been thoroughly explored. In contrast to its activity in adult tumors, SLFN11 did not predict favorable outcomes in pediatric patients, was not silenced, and promoted tumor growth. Resistance to replicative stress in SLFN11-expressing sarcomas was reversed by selective inhibition of BCL-XL.

## INTRODUCTION

Pediatric sarcomas are a heterogeneous group of malignancies disproportionately affecting adolescents and young adults. Multimodal therapy with chemotherapy, surgery, and radiation therapy (RT) has improved outcomes for those patients with localized disease. However, progress has stalled for patients with metastatic disease, who continue to have survival rates of <30% for the most common subtypes [1–5]. Therefore, novel therapeutic strategies and biomarkers that predict sensitivity to therapy are needed.

Previously, we reported that combining the poly (ADP-ribose) polymerase inhibitor (PARPi) talazoparib (TAL) with the topoisomerase I inhibitor (Topo1i) irinotecan (IRN) and temozolomide (TMZ) resulted in high rates of complete response (CR) in a murine model of Ewing sarcoma (ES) [6]. This work motivated a clinical trial testing TAL plus IRN with and without TMZ in children with refractory/recurrent solid tumors (NCT02392793). Results were encouraging: of 24 evaluable patients, 1 with Ewing sarcoma (ES) had a complete response (CR), 5 others had a partial response (PR), and 18 had disease stabilization [7]. However, it remains unclear how the combination works in this population.

Although BRCA mutations are rare in ES, several mechanisms have been proposed to explain the PARPi sensitivity in ES, most notably *functional* BRCA deficiency and SLFN11 [8–11]. Fusion proteins involving the peptide encoded by *EWSR1* at the N-terminus are the oncogenic drivers in ES (which has *EWSR1-FLI1/ERG* fusion) and in a subset of aggressive sarcomas, such as desmoplastic small round cell tumors (DSRCTs) (which have *EWSR1-WT1* fusion) and clear cell sarcomas (which have *EWSR1-ATF1* fusion). Gorthi et al [11] reported that the *EWSR1* fusion protein increases R-Loop formation, which sequesters BRCA1, rendering the tumor cell BRCA deficient and susceptible to replicative stress. They speculated that BRCA deficiency might create a liability in all tumors possessing an *EWSR1*-translocation. Tumors deficient in mediators of homologous recombination (HR) such as *BRCA1* and *BRCA2* are more susceptible to DNA single strand breaks: consequently, PARPis are selectively lethal in these cells. However, in contrast to PARPi treatment of adult tumors with *BRCA1* or *BRCA2* mutations, single-agent PARPi treatment in patients with relapsed/refractory ES elicited no significant responses or durable disease control [12].

Ewing sarcomas express high levels of Schlafen family member 11 (SLFN11), a putative DNA/RNA helicase whose expression has been associated with the response to DNA damaging agents (DDAs), thymocyte maturation, viral immunity, and interferon production [13]. SLFN11 augments sensitivity to replicative stress by stalling replication forks and impairing the DNA repair checkpoint response [14, 15]. PARPis cause replicative stress through PARP trapping [9], whereby the PARP protein becomes physically associated with DNA. This is similar to the mechanism of action of Topo1is, which are well-known inducers of replicative stress [16]. Additionally, PARP inhibition augments Topo1i toxicity by preventing the recruitment of repair enzymes to the site of damage, and TMZ enhances PARPi-mediated replicative stress by augmenting PARP trapping [17]. Therefore, the strategy of combining a PARPi, a Topo1i, and TMZ is a rational means of exploiting replicative stress in cancer cells. In retrospective studies, patients with ovarian cancer [18], breast cancer [19], prostate cancer [20], and ES-family tumors [10] who were treated with DDAs had better prognosis if their tumors had high SLFN11 expression. Patients with small-cell lung cancer expressing SLFN11 showed improved progression-free survival (PFS) and overall survival (OS) when treated with the PARPi veliparib and TMZ [21].

Determining whether *SLFN11* or the *EWSR1* fusion drives sensitivity to PARPi combinations in pediatric sarcomas is crucial for identifying patients who might benefit the most from such treatment. Although SLFN11 is highly expressed in ES, its expression pattern in other pediatric sarcomas is largely unknown. *SLFN11* mutations are rare, and epigenetic regulation has been suggested as a mediator of resistance [16, 22]. In this work, we show that SLFN11 is widely expressed in common pediatric sarcoma subtypes and that the SLFN11 protein, not the presence of a *EWSR1* fusion, drives sensitivity to TAL and IRN both *in vitro* and *in vivo*. Importantly, we show that SLFN11 expression does not portend a better prognosis in these patients: in fact, we reveal an unexpected oncogenic role for the protein. We also show that impairment of intrinsic apoptosis, not loss of SLFN11 expression, is a primary means of resistance to PARPi combination therapy in pediatric sarcoma, and that sensitivity to TAL+IRN can be restored by selective inhibition of BCL-XL. Our work supports the use of combinations involving strong-trapping PARPis and Topoi1s as targeted therapy for SLFN11-positive pediatric sarcomas, and it offers novel strategies to combat tumors resistant to replicative stress.

## RESULTS

### SLFN11 Expression Is Highly Correlated with Sensitivity to SN-38 and TAL

To determine the extent to which SLFN11 expression is correlated with sensitivity to DDA, we analyzed the Genomics of Drug Sensitivity in Cancer (GDSC) database which contains more than 1000 cell lines that have been assayed for cell viability 72 h after exposure to hundreds of drugs [23]. The efficacies of several DDAs, as measured by the area under the curve (AUC), were highly correlated with SLFN11 expression, with those of the strong-trapping PARPi TAL, and SN-38, the active metabolite of IRN, showing the highest statistical significance and the largest effect sizes (**Fig. 1A**; **Supplementary Table S1A**). In contrast, the microtubule inhibitors vinorelbine and vinblastine and the weak-trapping PARPi olaparib showed poor associations. The Pearson correlation between the mean AUC of SN-38 and TAL was 0.51 (*P* < 0.001), and there was a clear trend between this average and the increasing quintiles of SLFN11 expression (*P* < 0.001, 1-way ANOVA) (**Fig. 1B**; **Supplementary Table S1B**). We observed no correlation between drug response and those genetic lesions known to impair HR and sensitize tumor cells to both PARPis and Topo1is, such as BRCA1, BRCA2, and ATM (**Fig. 1C**; **Supplementary Fig. S1A and S1B**; **Supplementary Table S1C-S1E**) [24–26]. The short timescale of the GDSC viability assay (72 h) suggests that SLFN11 induces rapid cytotoxicity, and this phenotype appears to be distinct from that induced by HR defects.

**Figure 1.**
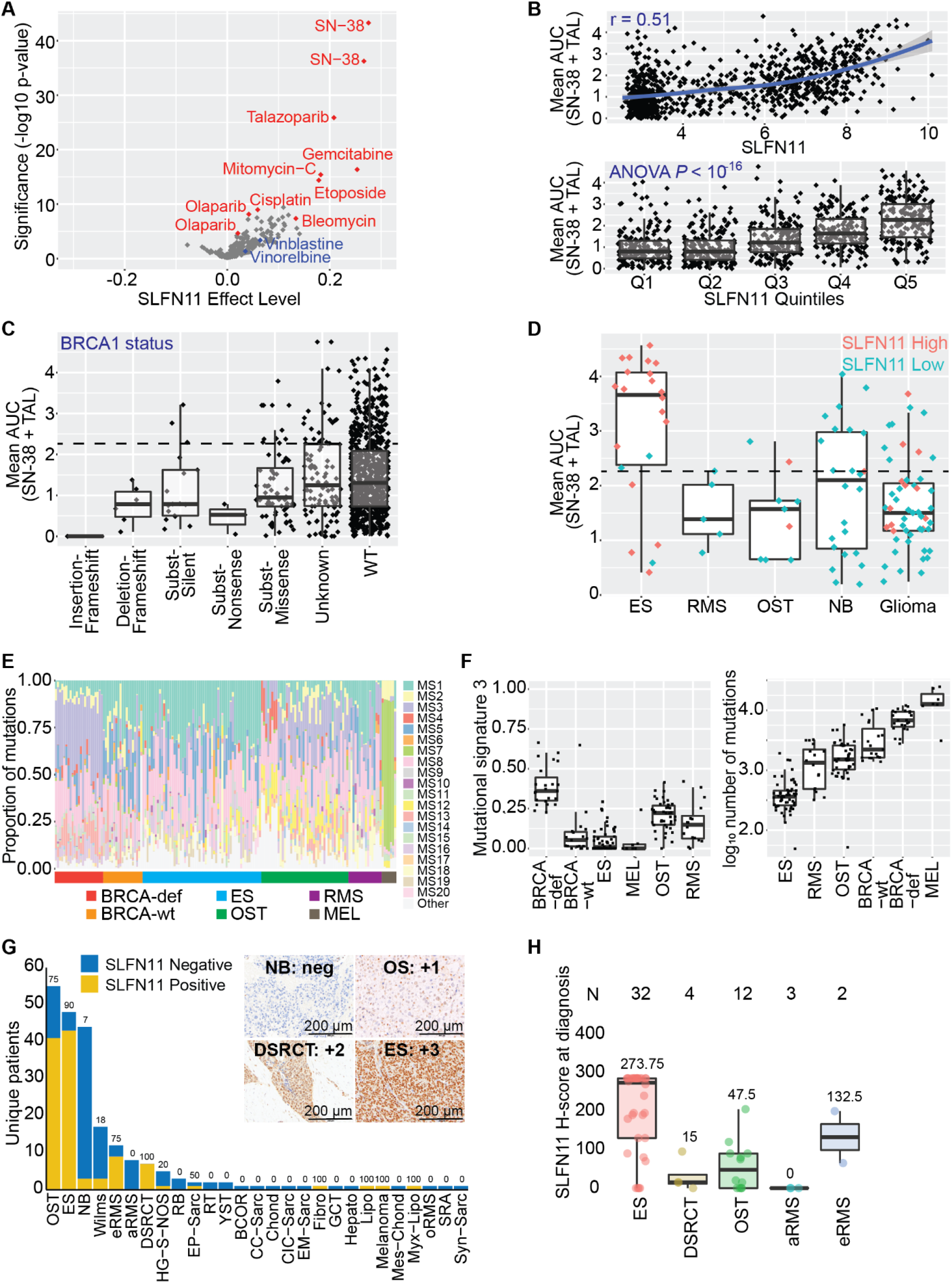
SLFN11 was highly correlated with the efficacy of SN-38 and TAL, and was widely expressed in pediatric sarcoma. Correlation between SLFN11 expression and the single-agent AUC **(A)** or mean **(B)** of the AUC for SN-38 and TAL after 72 h drug exposure as reported in the GDSC database. Olaparib and SN-38 appeared as two different batches in the database. **(C)** Correlation between the mean of the AUC for SN-38 and TAL from the GDSC and BRCA1 mutational status as annotated in the COSMIC database. Dotted line equals the median activity of the highest quintile of SLFN11 expression (“Q5”) from **(B)**. **(D)** Distribution of the mean of the AUC for SN-38 and TAL from the GDSC in Ewing sarcoma (ES), rhabdomyosarcoma (RMS), OST (osteosarcoma), NB (neuroblastoma), and glioma cell lines. Cells marked “SLFN11 High” (salmon) expressed the highest quintile of SLFN11 expression (“Q5”) from **(B),** while all others were defined as “SLFN11 Low” (teal). Dotted line equals the median activity of the highest quintile of SLFN11 expression (“Q5”) from **(B)**. **(E-F)** Mutational signatures and total number of mutations calculated from ES, RMS, and OST tumor samples from pediatric patients. BRCA-wt and BRCA-deficient samples were included as controls for MS3. Melanoma was included as a positive control for MS7. **(G)** SLFN11 status as assessed by immunohistochemistry in 353 samples from 220 unique patients treated on solid tumor protocols at our institution. “SLFN11 Negative” was defined as H-score = 0, and “SLFN11 Positive” as H-score > 0. **(H)** SLFN11 H-score at diagnosis for select pediatric sarcoma. The mean H-score is reported for each tumor type.

We further explored the GDSC database by first defining cell lines within the top quintile of expression as “SLFN11 High” and all others as “SLFN11 Low” then examining the distribution of AUCs by tumor subtype (**Fig. 1D**). The correlation between SLFN11 expression and average SN-38/TAL activity was present in ES, rhabdomyosarcoma (RMS), and osteosarcoma (OST) cell lines, although sampling was low for the latter 2 tumor types. In contrast, all but 1 neuroblastoma model expressed low SLFN11, despite 11/24 cell lines (46%) showing a drug response comparable to that of the highest quintile of SLFN11 expressors. Moreover, there was little difference in the drug response of high and low SLFN11 expressors in glioma cell lines, suggesting that the potential for SLFN11 to act as a biomarker predicting sensitivity to SN-38/TAL varies considerably across tumor types.

To better understand the relation between SLFN11 levels and DNA repair defects, we calculated mutational signatures (MS) in tumors from pediatric patients with ES, RMS, and OST (**Fig. 1E** **and** **1F**; **Supplementary Table S1F**), using data from BRCA-deficient and BRCA-wild-type cohorts used as control [27]. As expected, expression of MS3, a signature associated with HR repair defects, was highest in the BRCA-deficient group. Consistent with recent reports, OST also showed elevated expression of this signature [28]. In contrast, ES tumors had MS3 levels comparable to those in BRCA-wild-type tumors, a finding inconsistent with reports that suggest translocations involving *EWSR1* induce functional BRCA deficiency. Consistent with previously reported whole-genome sequencing studies [29, 30], ES tumors have low genomic instability as assessed by the total number of mutations—an observation incompatible with the presence of HR deficiency.

Given the high level of SLFN11 expression in ES and the relative scarcity of data on its expression in other pediatric sarcoma, we developed an immunohistochemistry (IHC) protocol that uses a commercially available antibody to assess protein levels in pediatric sarcomas directly. We assayed 353 samples from 220 different patients with non-CNS solid tumors who had sufficient material for staining at St. Jude Children’s Research Hospital (**Fig. 1G**; **Supplementary Table S1G and S1H).** The patient demographics and diagnosis groups are shown in **Table 1**. SLFN11 had variable expression, but was nearly universal in ES and DSRCT, with 90% and 100% of those tumors showing SLFN11 positivity (H-score > 0), respectively. SLFN11 was detected in 75% of the samples from patients with OST or embryonal RMS (eRMS). Quantification of SLFN11 in the other tumor types was limited by the small sample sizes. Using H-score at diagnosis, we found the highest SLFN11 expression in ES, followed by eRMS, OST, and DSRCT **(Fig. 1H**), with a few samples of the latter 3 tumor types having high expression levels similar to those observed in ES. Overall, SLFN11 was expressed in 69% of pediatric sarcoma sampled, and 76% of the most common pediatric sarcomas—a significantly higher percentage than has been implicated in adult tumors [31].

**Table 1.**
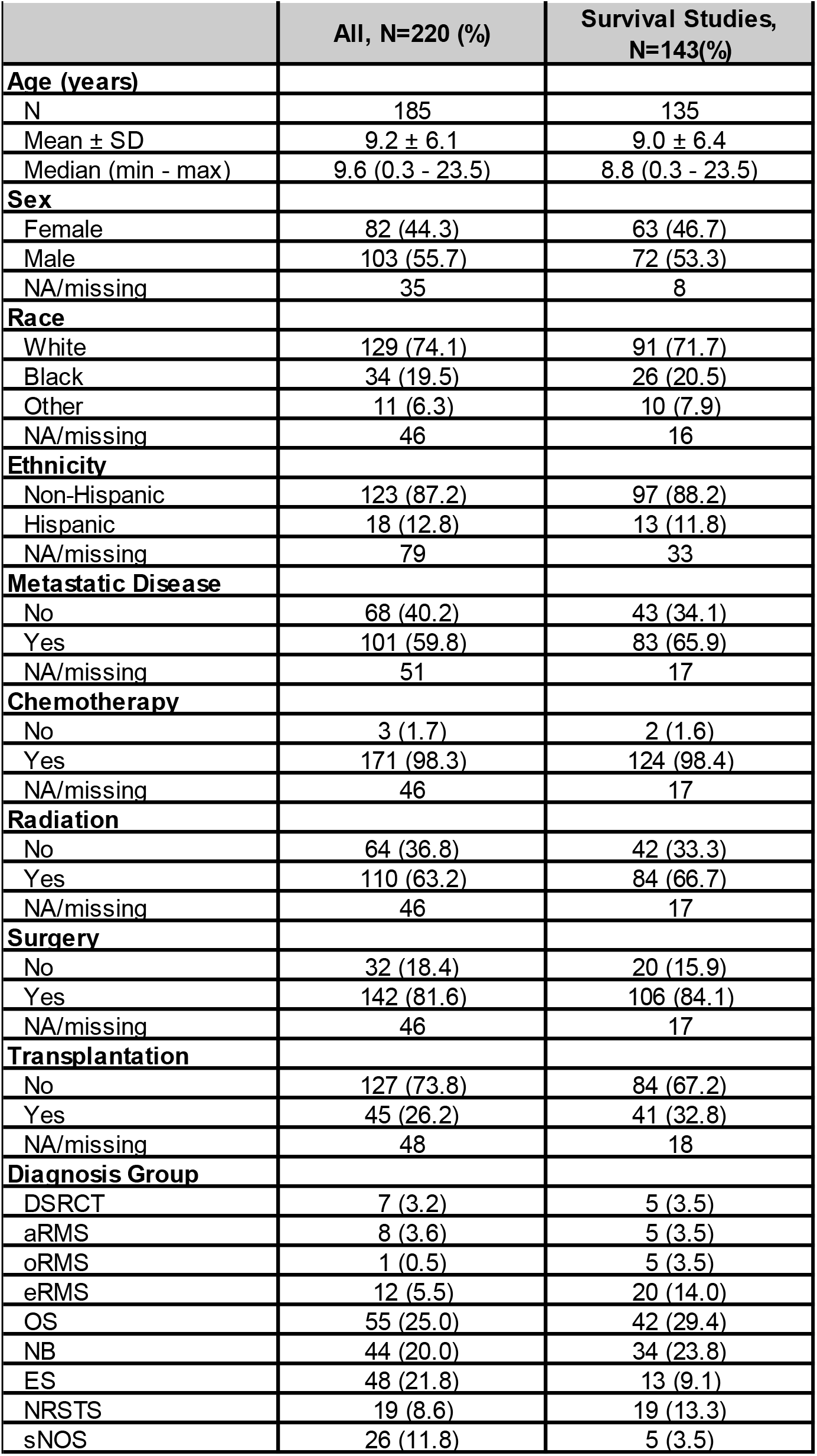
Demographics of thepediatric cohort.

|  | All, N=220 (%) | Survival Studies, N=143(%) |
| --- | --- | --- |
| **Age (years)** |  |  |
| N | 185 | 135 |
| Mean ± SD | 9.2 ± 6.1 | 9.0 ± 6.4 |
| Median (min - max) | 9.6 (0.3 - 23.5) | 8.8 (0.3 - 23.5) |
| **Sex** |  |  |
| Female | 82 (44.3) | 63 (46.7) |
| Male | 103 (55.7) | 72 (53.3) |
| NA/missing | 35 | 8 |
| **Race** |  |  |
| White | 129 (74.1) | 91 (71.7) |
| Black | 34 (19.5) | 26 (20.5) |
| Other | 11 (6.3) | 10 (7.9) |
| NA/missing | 46 | 16 |
| **Ethnicity** |  |  |
| Non-Hispanic | 123 (87.2) | 97 (88.2) |
| Hispanic | 18 (12.8) | 13 (11.8) |
| NA/missing | 79 | 33 |
| **Metastatic Disease** |  |  |
| No | 68 (40.2) | 43 (34.1) |
| Yes | 101 (59.8) | 83 (65.9) |
| NA/missing | 51 | 17 |
| **Chemotherapy** |  |  |
| No | 3 (1.7) | 2 (1.6) |
| Yes | 171 (98.3) | 124 (98.4) |
| NA/missing | 46 | 17 |
| **Radiation** |  |  |
| No | 64 (36.8) | 42 (33.3) |
| Yes | 110 (63.2) | 84 (66.7) |
| NA/missing | 46 | 17 |
| **Surgery** |  |  |
| No | 32 (18.4) | 20 (15.9) |
| Yes | 142 (81.6) | 106 (84.1) |
| NA/missing | 46 | 17 |
| **Transplantation** |  |  |
| No | 127 (73.8) | 84 (67.2) |
| Yes | 45 (26.2) | 41 (32.8) |
| NA/missing | 48 | 18 |
| **Diagnosis Group** |  |  |
| DSRCT | 7 (3.2) | 5 (3.5) |
| aRMS | 8 (3.6) | 5 (3.5) |
| oRMS | 1 (0.5) | 5 (3.5) |
| eRMS | 12 (5.5) | 20 (14.0) |
| OS | 55 (25.0) | 42 (29.4) |
| NB | 44 (20.0) | 34 (23.8) |
| ES | 48 (21.8) | 13 (9.1) |
| NRSTS | 19 (8.6) | 19 (13.3) |
| sNOS | 26 (11.8) | 5 (3.5) |

### SLFN11, Not *EWSR1* Translocation, Is Required for Sensitivity to SN-38 and TAL *In vitro*

To further assess SLFN11 and *EWSR1* translocation as potential drivers of sensitivity to TAL and SN-38, we profiled 14 sarcoma cell lines that varied by translocation type, p53 status, and histology, then we assessed SLFN11 expression by IHC, Western blot, and qPCR analysis (**Fig. 2A**, **Supplementary Table S2A and S2B**). The Pearson correlation between SLFN11 protein and mRNA levels was 0.64 (*P* = 0.018) (**Supplementary Fig. S2A**). Protein levels were correlated with sensitivity to single-agent SN-38 and TAL, with Pearson correlations of 0.72 (*P* = 0.003) and 0.74 (*P* = 0.003), respectively (**Supplementary Fig. S2B**); and with the mean AUC of the 2 compounds, with a Pearson correlation of 0.77 (*P* = 0.001) (**Fig. 2B)**. We found no association between sensitivity to these drugs and p53 status (*P* = 0.12, *t-test*) (**Supplementary Fig. S2C**). Although most *EWSR1-*translocated cell lines were more sensitive to drug treatment when compared to non-translocated cell lines, they also tended to express the highest levels of SLFN11. The exception was SU-CCS-1, a SLFN11-negative (no protein detected by Western; IHC H-score = 0) *EWSR1-ATF1*-translocated clear cell sarcoma, suggesting that the *EWSR1* translocation alone was insufficient to drive drug sensitivity.

**Figure 2.**
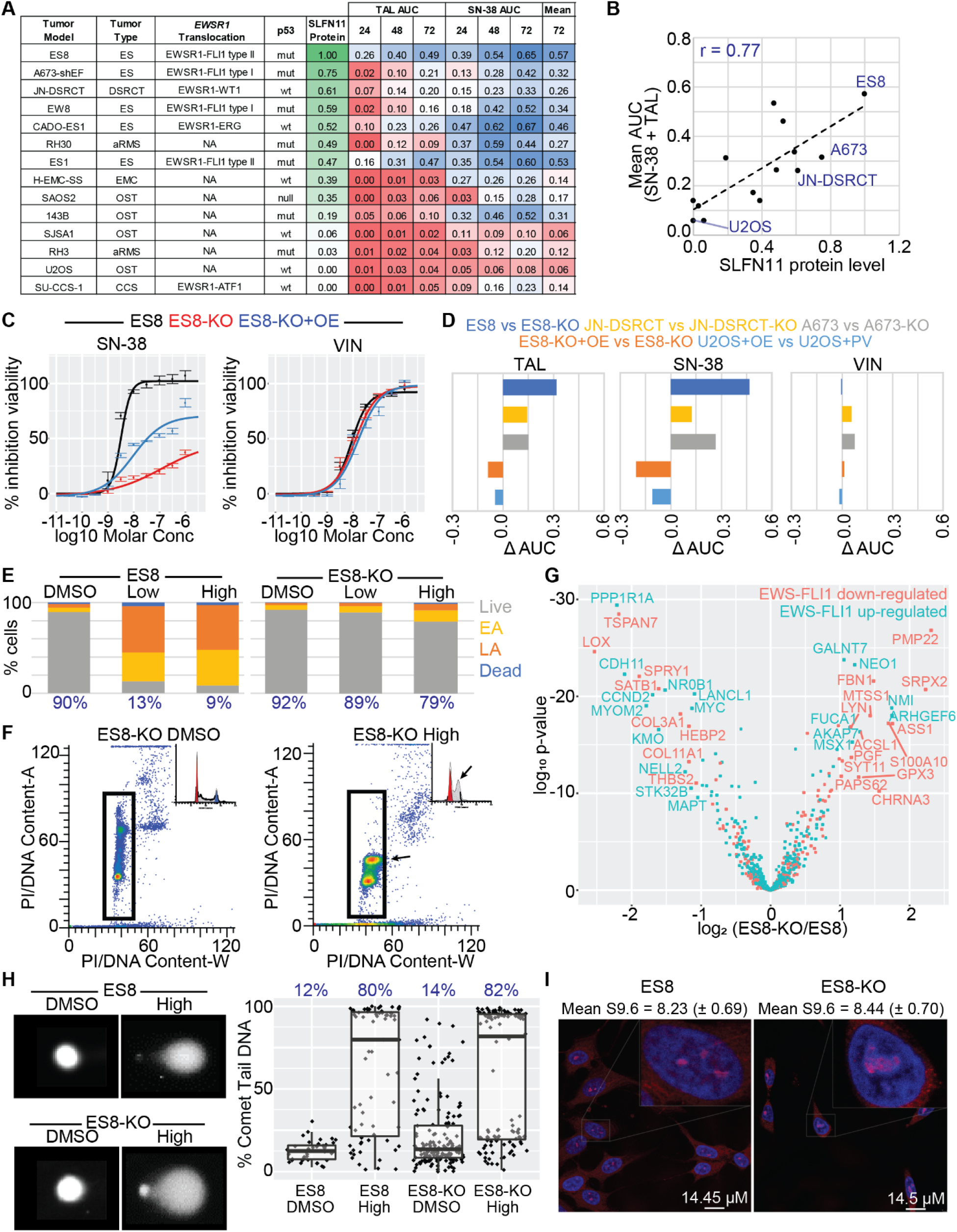
SLFN11, not *EWSR1*-translocation or R-loop expression, predicted sensitivity to SN-38 and TAL *in vitro*. **(A)** SLFN11 protein levels and CellTiter-Glo (CTG) results for 14 sarcoma cell lines that varied by translocation status, p53 status, and histology. Protein levels were normalized to ES8. The area-under-the-curve (AUC) for the dose-response curve was calculated at 24, 48, and 72 hours in the concentration range 10^−11^ to 10^−4^ Molar. Each value was then normalized to 700 – the maximum observed AUC for 100% efficacy at all concentrations in the range – yielding a number from 0 to 1. Although extraskeletal myxoid chondrosarcoma (EMC) cancers such as H-EMC-SS typically have *EWSR1-NR4A3* fusions, we did not detect a *EWSR1*-translocation in this line. n ≥ 2. **(B)** Correlation between SLFN11 protein levels and the mean of the AUC for SN-38 and TAL for the cell panel in **(A)**. **(C)** CTG dose-response following 72h drug exposure of SN-38 and vincristine in ES8, ES8-KO, and ES8-KO+OE cells. n ≥ 2. **(D)** Difference in normalized CTG AUC following 72 h drug exposure of TAL, SN-38, and vincristine in knockout and over-expression models. n ≥ 2. **(E)** Flow cytometry assessment of cytotoxicity in ES8 and ES8-KO cells following 24h exposure to ‘Low’ (10 nM SN-38 + 10 nM TAL) and ‘High’ (1 μM SN-38 and 1 μM TAL) drug combinations. The percent live cells is reported in blue. EA = early apoptosis. LA = late apoptosis. n ≥ 2. **(F)** Cell cycle analysis of ES8-KO cells following 24 h exposure to ‘Low’ SN-38+TAL. Arrows highlight the build-up of S-phase cells induced by the drug combination. **(G)** Volcano plot showing the difference in expression of reported EWS-FLI1 downregulated (salmon) and upregulated (teal) genes between ES8 and ES8-KO cells. Expression was assessed by microarray at 4 h and 24 h following exposure to 0 and 2 Gy. n = 3. **(H)** Exemplar image and quantification of the alkaline comet tail assay of ES8 and ES8-KO cells following 2.5h exposure to the ‘High’ concentration of SN-38 and TAL. Mean percent comet tail DNA is reported in blue. **(I)** Immunofluorescence quantification of R-loops in untreated ES8 and ES8-KO cells. Nuclei were stained with DAPI (blue) and R-loops were stained with S9.6 antibody (red). Mean (sem) S9.6 nuclear intensity is reported.

To confirm SLFN11 as the primary driver of sensitivity to these agents, we knocked out the gene by using CRISPR/Cas9 in our 3 highest-expressing models, generating the isogenic pairs: ES8/ES8-KO, JN-DSRCT/JN-DSRCT-KO, and A673/A673-KO. We also over-expressed the protein in U2OS cells, which showed little baseline expression, to create U2OS-OE cells (61% protein relative to total ES8 by Western); and in ES8-KO cells, to create ES8-KO+OE cells (37% protein relative to total ES8). Knockout and over-expression were confirmed by Western blot analysis and IHC (**Supplementary Figure S2D and S2E**; **Supplementary Table S2A and S2B**). Loss of SLFN11 protein significantly reduced sensitivity to both SN-38 and TAL in all three knockout lines, whereas over-expression in U2OS and ES8-KO cells increased drug sensitivity (**Fig. 2C** **and** **2D**). Consistent with our GDSC analysis, SLFN11 loss had little effect on vincristine sensitivity. Despite expressing a lower level of the same engineered SLFN11 protein construct, ES ES8-KO+OE cells had higher AUC values for SN-38 and TAL when compared to OST U2OS-OE cells (0.40 vs. 0.25 for SN-38 and 0.26 vs. 0.10 for TAL), consistent with the hypothesis that the magnitude of sensitization induced by SLFN11 varies between tumor types (**Supplementary Fig. S2F**). Ionizing radiation is another means to induce replicative stress [32]. In agreement with our SN-38 and TAL experiments, ES8-KO and JN-DSRCT-KO were more viable, and U2OS-OE was less viable, compared to their wild-type counterparts at 72 h after exposure to 4 Gy radiation (**Supplementary Fig. S2G**).

To further study the effect of SLFN11 KO in our models, we used flow cytometry to compare cell cycle effects and the degree of cell death induced by “Low” (10 nM SN-38 + 10 nM TAL) and “High” (1000 nM SN-38 + 1000 nM TAL) dose combinations after 24 h exposure. Based on our previous pharmacokinetic assessment, “Low” approximates to the upper bound of clinically relevant concentrations for both drugs, whereas “High” is physiologically unobtainable but useful for studying mechanism and resistance [6]. Wild-type ES8 cells showed near-complete loss of viability at both “Low” and “High” doses of the combination, whereas the ability of SN-38+TAL to induce cell death in ES8-KO cells was significantly diminished, even at high concentrations (**Fig. 2E)**. A similar decrease in cell viability was observed in JN-DSRCT compared to JN-DSRCT-KO cells, and the combination was also less cytotoxic in SLFN11-negative SU-CCS-1 cells (**Supplementary Fig. S2H**). SLFN11 selectively induces death in cells arrested in S-phase because of replicative stress [15]. Consistent with this finding, we observed a significant build-up of S-phase-arrested ES8-KO cells (**Fig. 2F**).

Previous studies have shown that SLFN11 is a transcriptional target of EWS-FLI1 [10]. To determine whether SLFN11 itself contributes to the regulation of EWSR1-FLI1 target genes, we assessed gene expression in ES8 and ES8-KO cells at 4 h and 24 h following exposure to 0 or 2 Gy of ionizing radiation (baseline and stress conditions). EWSR1-FLI known targets mapped equally between upregulated and downregulated genes, and we found no enrichment of directional activation or inhibition of those EWSR1-FLI1 target genes when using Gene Set Enrichment Analysis (GSEA, FDR > 0.05) (**Fig. 2G**; **Supplementary Table S2C)**. Therefore, although SLFN11 is regulated by EWS-FLI1, it appears to perturb gene expression independently of the fusion protein.

To determine whether SLFN11 influenced the extent of DNA damage induced by SN-38 and TAL, we performed an alkaline comet tail assay after exposing ES8 and ES8-KO cells to DMSO and the “High” concentration of SN-38 and TAL for 2 h (**Fig. 2H**). The amount of DNA damage was similar in both cell lines after drug treatment, indicating that SLFN11 does not enhance the degree of damage but rather increases the probability of cell death following drug insult. Finally, given the findings of high levels of R-loops in *EWSR1*-translocated tumors [11], we quantified R-loop expression in ES8 and ES8-KO cells. Despite a remarkable difference in their response to SN-38 and TAL, we found no significant difference in R-loop levels in the wild-type and SLFN11 KO models (**Fig. 2I)**.

### SLFN11, Not *EWSR1* Translocation, Is required for Sensitivity to TAL and IRN *In Vivo*

To confirm our *in vitro* finding indicating that SLFN11 was an important driver of drug response in *EWSR1*-translocated tumors, we conducted *in vivo* efficacy studies, as described previously [6], using luciferase-labeled xenografts of ES8, ES8-KO, JN-DSRCT, JN-DSRCT-KO, and SU-CCS-1 cells. Mice were screened weekly by Xenogen^®^ imaging and enrolled in the study after a target bioluminescence signal of 10^7^ photons/s/cm^2^ or a palpable tumor was obtained. We used clinically relevant doses and schedules for all treatment groups tested (**Fig. 3A**) [6] and administered 4 courses of therapy (21 days/course).

**Figure 3.**
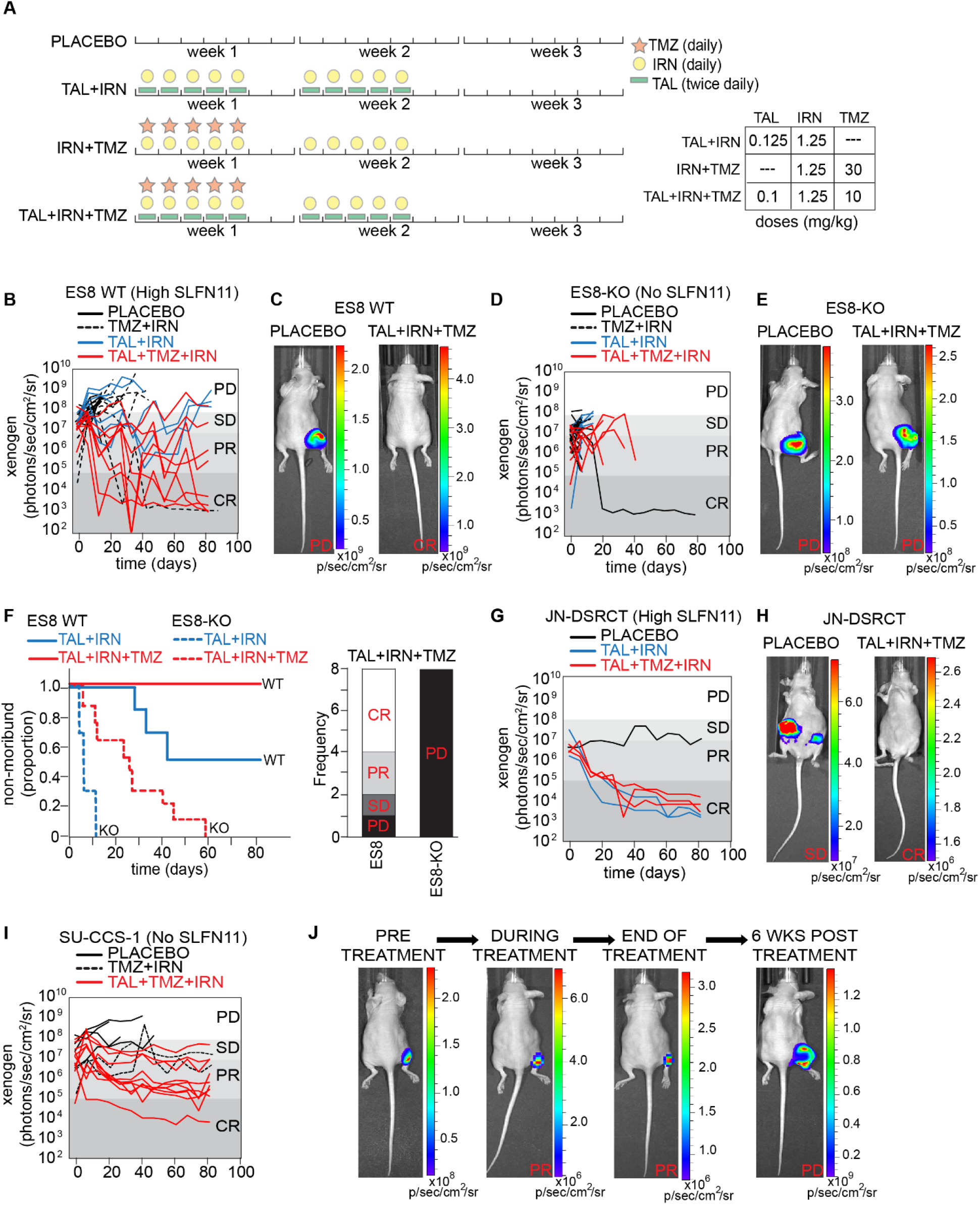
SLFN11, not *EWSR1*-translocation, was required for sensitivity to SN-38 and TAL *in vivo.* **(A)** Drug schedule selected for each combination treatment regimen. **(B)** Line plot of tumor burden over time as measured by bioluminescence for ES8 xenografts. Each line is a different mouse. **(C)** Representative images from the study in **(B)** are shown with PD in the placebo control group and a CR in the TAL+TMZ+IRN group. **(D)** Line plot of tumor burden over time for ES8-KO xenografts. **(E)** Representative images from the study in **(D)** are shown with PD in both the placebo and TAL+TMZ+IRN groups. **(F)** Survival curves and response at end of treatment with TAL+TMZ+IRN for ES8 and ES8-KO models. **(G)** Line plot of tumor burden over time for JN-DSRCT xenografts. **(H)** Representative images from the study in **(G)** are shown with PD in the placebo group and a CR in the TAL+TMZ+IRN group. **(I)** Line plot of tumor burden over time for SU-CCS-1 xenografts. **(J)** Representative images of a mouse treated over time in the TAL+TMZ+IRN group from **(I)** showing a PR during treatment and regrowth of tumor upon stopping therapy. PD, progressive disease; SD, stable disease; PR, partial response; CR, complete response.

In ES8 (high SLFN11) xenografts, mice treated with TAL+IRN+TMZ had the best response, with 100% surviving the 84 days of therapy, and 75% experiencing a CR or PR as determined by the bioluminescence signal (**Fig. 3B** **and** **3C**; **Supplementary Table S3**). Mice treated with TAL+IRN survived an average of 60.5 days, with 50% surviving all 4 courses of therapy. In sharp contrast to mice with wild-type ES8 xenografts, mice with ES8-KO xenografts treated with TAL+IRN and TAL+IRN+TMZ survived an average of 7.5 days and 23.1 days, respectively, with 100% having progressive disease (PD) (**Fig. 3D** **and** **3E**; **Supplementary Table S3**). No mice in any ES8-KO treatment cohort survived all 4 courses of therapy, although 1 control mouse appeared to have lost its engraftment signal after 2 weeks and survived the study. Compared to ES8 mice, survival in ES8-KO mice was significantly lower when treated with either TAL+IRN (*P* < 0.001) or TAL+IRN+TMZ (*P* < 0.001) (**Fig. 3F**).

Testing JN-DSRCT (high SLFN11) xenografts *in vivo* was challenging, as these tumors grew slower and had lower engraftment rates by comparison with ES8 xenografts. Five wild-type mice were successfully enrolled and treated with either TAL+IRN or TAL+IRN+TMZ, and all experienced a CR by 84 days (**Fig. 3G** **and** **3H**; **Supplementary Table S3**). One untreated mouse was also enrolled and maintained stable disease (SD) throughout the 4 courses of therapy. Interestingly, JN-DSRCT-KO cells engrafted poorly and were unable to be tested. In SU-CCS-1 (no SLFN11) xenografts, disease stabilized in mice treated with TAL+IRN+TMZ and 90% of the mice had SD or a PR at the end of therapy (**Fig. 3I**). However, once therapy was stopped, all mice regrew tumors within a few weeks (**Fig. 3J**). Together, these findings confirm the importance of SLFN11 in driving *in vivo* sensitivity to combinations involving SN-38 and TAL in *EWSR1*-translocated tumors. The most striking result was the near complete loss of efficacy in ES8-KO xenografts.

Finally, we explored efficacy in 143B cells, an aggressive OST cell line (*EWSR1* fusion negative) with low SLFN11 expression (19% of that in ES8 cells), and observed an intermediate response. Mice treated with TAL+IRN survived an average of 13.4 days, with 100% having PD (**Supplementary Fig. S3**; **Supplementary Table S3**). However, mice treated with TAL+IRN+TMZ survived much longer, with an average of 77.8 days on study, although none experienced a CR.

### SLFN11 Positivity Is Not Associated With Better Outcomes in Children With Sarcoma

Motivated by the strong evidence that SLFN11 sensitized pediatric sarcomas to PARPi combination therapy *in vitro* and *in vivo*, we performed a retrospective analysis of the patient cohort profiled in our IHC study to determine how SLFN11 status changed as therapy progressed and whether protein levels predicted clinical outcome. Only patients who had a sample available prior to recurrence or progression were included in the survival analysis (N = 143, **Table 1**). 98.4% of patients were treated with at least one DDA and 66.7% received radiation at some point in their therapy. This population was more refractory than would be expected historically, with patients with NRSTS, ES, and eRMS all having OS and event-free survival (EFS) of <50% at 5 years (**Fig. 4A**; **Supplementary Fig. S4A).**

**Figure 4.**
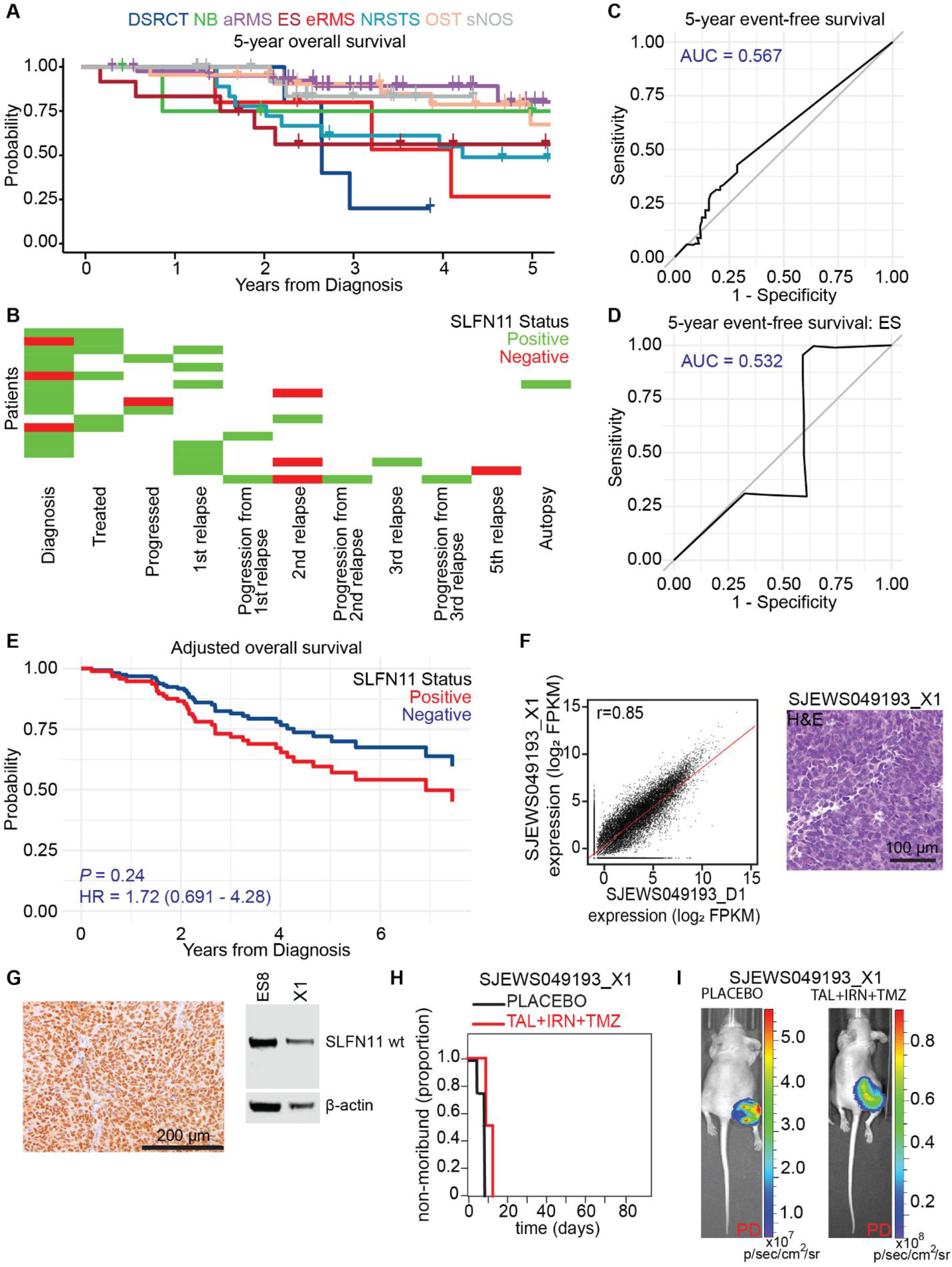
SLFN11 expression was not associated with better outcomes in children with sarcomas. **(A)** Five-year overall survival rates by diagnosis for the sarcoma patients in our IHC study. **(B)** SLFN11 status for 18 patients with at least two SLFN11 IHC measurements spanning different points in treatment. “Negative” and “Positive” SLFN11 status were defined as H-score equal to zero and H-score greater than 0, respectively. **(C)** ROC curve for five-year event free survival of all sarcoma patients as a function of H-score. **(D)** ROC curve for five-year event free survival of ES patients as a function of H-score. **(E)** Adjusted overall survival as a function of SLFN11 status after controlling for age, metastatic status, and disease. **(F)** RNA-seq expression profile comparing the ES patient-derived orthotopic xenograft model SJEWS049193_X1 and the matched primary tumor (SJEWS049193_D1, Pearson *r* = 0.85), and hematoxylin and eosin stain of SJEWS049193_X1. **(G)** SLFN11 IHC and Western blot from a SJEWS049193_X1 tumor sample. **(H)** Survival curves for SJEWS049193_X1 and **(I)** representative images from the efficacy study showing progressive disease (PD) in both the placebo and TAL+TMZ+IRN groups.

Loss of SLFN11 expression has been proposed as a mechanism of DDA resistance in adult tumors [33]. To explore the trend of SLFN11 levels in our cohort, we identified 18 patients with at least 2 IHC measurements spanning different points in treatment (**Fig. 4B**; **Supplementary Fig. S4B**). We found no evidence that SLFN11 was silenced over time: 10/18 of patients remained positive throughout, 3/18 went from being SLFN11 negative to SLFN11 positive, 3/18 went from being SLFN11 positive to SLFN11 negative, and 2/18 were initially SLFN11 positive then had a negative sample followed by another positive sample. When including all patients in our IHC study, 37.2% of those with a sample at diagnosis were positive, compared with 46.2% at progression and 53.7% at relapse (**Supplementary Table S4A**). We observed no clear trend in the mean values for SLFN11 H-score in samples obtained at diagnosis, relapse, or autopsy.

Using receiver-operating characteristic (ROC) analysis, we found that SLFN11 expression as measured by H-score failed to discriminate patients by EFS, OS, recurrence-free survival (RFS), or PFS across all tumor types (**Fig. 4C**; **Supplementary Fig. S4C-S4E**), or within individual disease cohorts (**Fig. 4D**; **Supplementary Fig. S4F and S4G**). Surprisingly, SLFN11 positivity was a statistically significant predictor of *worse* outcome in terms of RFS (*P* = 0.045, HR = 1.76 [1.01-3.05]) in univariate analysis (**Supplementary Table S4B**). The metrics OS (*P* = 0.072, HR = 1.82 [0.95-3.49]), EFS (*P* = 0.10, HR = 1.58 [0.92-2.73]), and PFS (*P* = 0.078, HR = 1.72 [0.94-3.13]) were nonsignificantly associated with poorer outcomes in patients with SLFN11 expressing tumors. Although statistical significance was not achieved for any metric in multivariate analysis controlling for age, metastatic status, and disease, the hazard ratio for positivity remained at or greater than unity (**Fig. 4E**; **Supplementary Fig. S4H-S4J).** ROC analysis using the H-scores failed to discriminate patients with metastatic disease (AUC = 0.605). Moreover, neither SLFN11 H-score nor positivity were statistically significant predictors of metastasis in multivariate analysis that controlled for diagnosis (*P* = 0.58 and 0.35, respectively).

To further interrogate the impact of SLFN11 expression in pediatric sarcoma, we developed and characterized an orthotopic xenograft model, SJEWS049193_X1, using a tumor obtained at autopsy from a patient with metastatic ES treated with multiple salvage regimens, including treatment on the TAL+IRN+TMZ clinical trial NCT02392793 (**Fig. 4F)**. Despite expressing a high level of SLFN11 (H-score = 285), this tumor failed to respond to TAL+IRN+TMZ *in vivo*, and 100% of mice showed PD (**Fig. 4G-4I**, **Supplementary Table S3**). Taken together, our clinical findings indicated that (a) SLFN11 positivity is common in pediatric sarcoma; (b) pediatric sarcomas do not acquire DDA resistance via *SLFN11* silencing; and (c) SLFN11 positivity does not predict improved patient outcomes, and might, in fact, do the opposite. These results suggest an oncogenic role for SLFN11 in these tumors.

### SLFN11 Knockout Induces Widespread Transcriptional Changes in Models of ES and DSRCT

To investigate the effect of SLFN11 beyond its role as a sensitizer to DNA damage, we expanded the microarray study described earlier to include JN-DSRCT and JN-DSRCT-KO cells (**Fig. 5A**). After controlling for cell of origin, 4960/21148 of the genes studied (23%) were differentially expressed in ES8 and JN-DSRCT upon SLFN11 loss (ANOVA FDR < 0.05). GSEA using the MsigDB Hallmark sets [34] revealed significant upregulation of genes involved in “Interferon Alpha Response”, “Interferon Gamma Response”, and “TNFA Signaling via NFKB”; and downregulation of genes enriched in “E2F targets” and “G2M Checkpoint” (FDR q-value < 0.001, **Fig. 5B**; **Supplementary Fig. S5A**) in SLFN11-KO cells. Ingenuity Pathway Analysis (IPA) [35], based on differentially expressed genes showing concordant change greater than 0.2 log in both cell backgrounds, confirmed “Interferon Signaling” activation in SLFN11-KO cells. The top regulator effect identified by IPA was enhanced “Quantity of MHC Class I on cell surface” in SLFN11-KO cells (**Fig. 5C**). This expression study suggests that loss of SLFN11 decreases proliferation and reduces evasion of the host innate immune response in these models.

**Figure 5.**
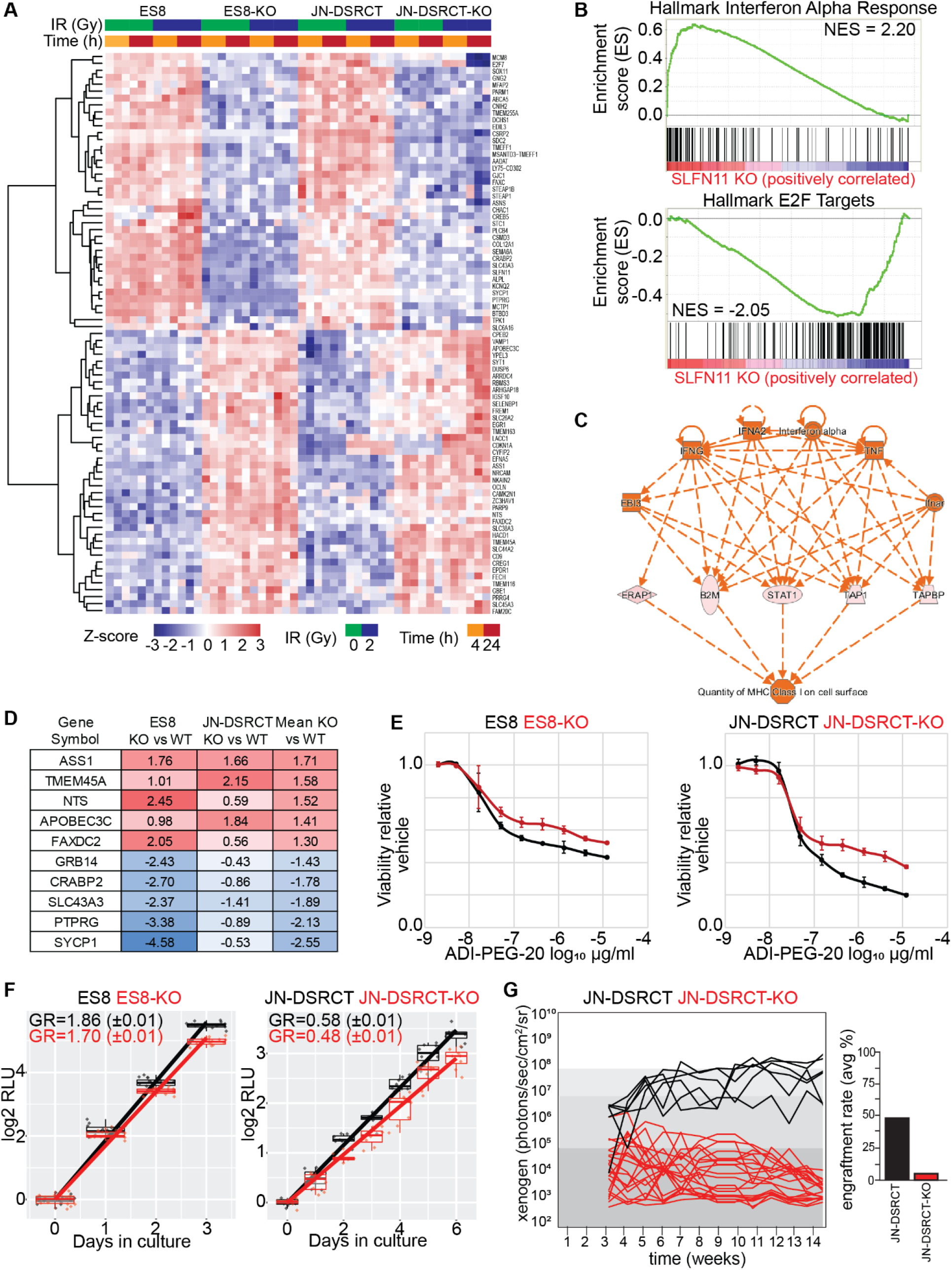
SLFN11 augmented metabolism, evasion from the innate immune response, and proliferation in pediatric sarcomas. **(A)** Heatmap of the most consistently differentially expressed genes (≥0.5 log_2_ unit change in the same direction) from a microarray experiment comparing ES8 and JN-DSRCT WT and KO cell lines at 4 h and 24 h following exposure to 0 and 2 Gy. n = 3. **(B)** Highest scoring GSEA Hallmark gene sets enriched in KO (top) and wild-type (bottom) lines. **(C)** The top regulator in KO cells identified via IPA upstream regulator analysis. **(D)** Top five most differentially activated or inhibited genes (log_2_ change in expression) in KO vs wild-type cells. **(E)** CTG assay assessing viability in KO and wild-type cells exposed to the pegylated arginine deiminase ADI-PEG-20. **(F)** Growth rate (GR) determination in KO vs wild-type cells, expressed as doublings/day (sem). **(G)** *In vivo* growth rate assessment comparing JN-DSRCT wild-type and KO models. Each cell line was injected at a cell density of 1 million cells/mouse. Xenogen measurements were started after 3-4 weeks for JN-DSRCT or JN-DSRCT-KO, then continued weekly.

The most upregulated gene in SLFN11-KO cells was that encoding arginase succinate synthetase 1 (ASS1), an enzyme involved in the arginine biosynthetic pathway (**Fig. 5D**). Others have shown that tumor cells repress the *ASS1* gene to bypass the urea cycle and increase cytosolic aspartate levels, thereby stimulating pyrimidine biosynthesis and promoting cancer proliferation [36, 37]. ASS1 deficiency is common in sarcomas, and renders them vulnerable to depletion of arginine from their environment via pegylated arginine deiminase (ADI-PEG 20) [38, 39]. To test whether changes in *ASS1* expression had functional consequences, we treated ES8 and JN-DSRCT wild-type and SLFN11-KO cells with ADI-PEG 20 in a dose-response experiment and then evaluated cell viability with the CellTiter-Glo (CTG) assay. Consistent with our expression study, wild-type cells were more sensitive to arginine depletion than were SLFN11-KO cells (**Fig. 5E**).

To determine whether the presence of SLFN11 conferred a general growth advantage to sarcoma cells, we measured the growth rate of wild-type and SLFN11-KO cells in culture. We observed decreases of 9%, 17%, and 27% in the growth rates for KO cells compared to wild-type ES8, JN-DSRCT, and A673 cells, respectively; and an increase of 7% for U2OS-OE cells compared with U2OS cells (**Figure 5F**; **Supplementary Fig. S5B**). *In vivo*, ES8 and ES8-KO models maintained the aggressive growth observed *in vitro* and we were unable to detect a difference in survival between untreated mice from these two groups (*P* = 0.316) (**Supplementary Table S3**). Strikingly, SLFN11 deletion caused a complete loss of the ability of JN-DSCRT-KO cells to form tumors in our xenograft models, despite multiple attempts to optimize conditions both intraperitoneally and subcutaneously (**Fig. 5G)**.

### Mechanisms of Resistance to SN-38 and TAL in SLFN11-Expressing Pediatric Sarcomas

The lack of SLFN11 silencing in the pediatric sarcoma population led us to seek alternative mechanisms of resistance in cell lines with high levels of SLFN11 expression. Unfortunately, SJEWS049193_X1, the resistant ES model described earlier, was not amenable to prolonged cell culture and *in vitro* investigation. Therefore, we mined the GDSC database to identify the sarcoma cell lines with the highest levels of SLFN11 expression that were refractory (having less than the median activity observed in the GDSC) to both SN-38 and TAL. We identified 3 lines of interest: EW-13, EW-18, and EW-24 (**Supplementary Fig. S6A**, bottom-left quadrant). These lines showed broad resistance to oncology drugs, including vincristine, when compared to sensitive ES models with high SLFN11 expression (**Fig. 6A**; **Supplementary Table S6A)**. We obtained EW-13 and EW-18, as well as EW-11 which showed intermediate sensitivity in the database. Separately, we acquired CHLA258, which was reported to be less sensitive to Topo1is [40]. We confirmed drug resistance with CTG assays and flow cytometry (**Fig. 6B**; **Supplementary Fig. S6B**). Western blot analysis indicated strong SLFN11 expression, with bands of appropriate size, in EW-11, EW-13, and CHLA258; however, protein expression in EW-18 was weaker and the band migration corresponded to a lower molecular weight (**Fig. 6C**; **Supplementary Fig. S6C**). Cross-referencing with the COSMIC database (https://cancer.sanger.ac.uk/cosmic) revealed that the *SLFN11* gene in EW-18 had a frameshift mutation, c.1928_1929insA, which resulted in a C-terminal truncation that was predicted to limit nuclear localization and, therefore, reduce DDA sensitization [14]. Indeed, IHC confirmed that SLFN11 was predominantly expressed in the cytoplasm, and consequently, the cell line was assigned an H-score of 0 (**Fig. 6D**; **Supplementary Fig. S6D**). It is important to note that, as far we are aware, all commercially available antibodies target the N-terminus of SLFN11 and could fail to detect C-terminal truncations unless a distinction is made between nuclear and cytoplasmic staining as we did in this study.

**Figure 6.**
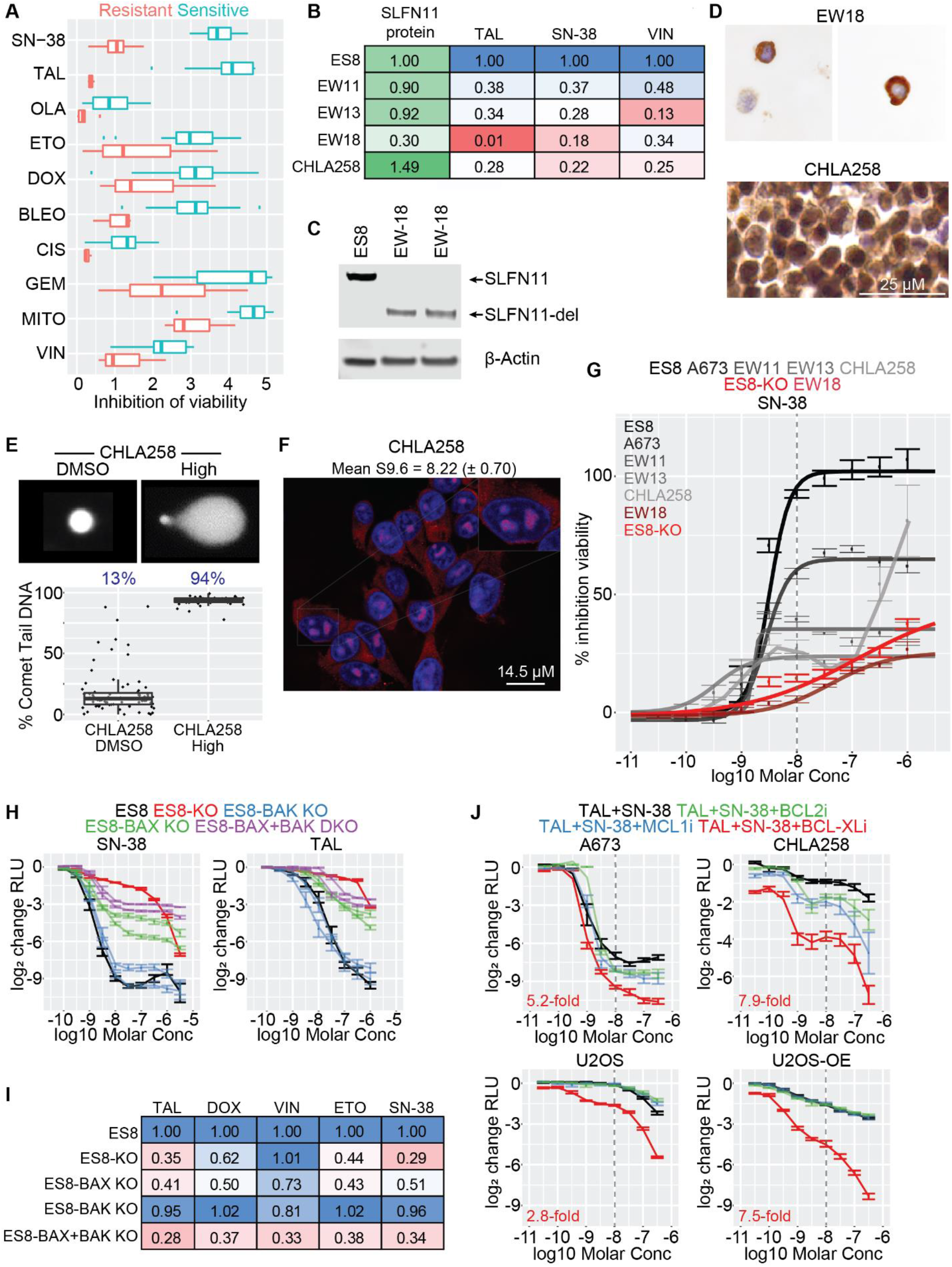
BCL-Xli restored sensitivity to SN-38 and TAL in resistant SLFN11 expressing sarcoma cells. **(A)** Boxplot comparing drug sensitivity in resistant and sensitive ES cell lines from the GDSC. **(B)** Heatmap of AUC values in four resistant ES cell lines normalized to ES8 (CTG assay, 72h drug exposure). n ≥ 2. **(C)** Western blot from two biological replicates of EW-18 confirming expression of a truncated SLFN11 protein. **(D)** Confirmation of cytoplasmic and nuclear staining of SLFN11 in EW-18 and CHLA258, respectively. **(E)** Exemplar image and quantification of the alkaline comet tail assay of CHLA258 cells following 2.5 h exposure to the “High” concentration of SN-38 and TAL. Mean percent comet tail DNA is reported in blue. **(F)** Immunofluorescence quantification of R-loops in untreated CHLA258 cells. Nuclei were stained with DAPI (blue) and R-loops were stained with S9.6 antibody (red). Mean (sem) S9.6 nuclear intensity is reported. **(G)** Dose-response curves for SN-38 (CTG, 72 h) in ES8 (black), SLFN11 expressing resistant ES cell lines (grays), ES8 KO (red), and EW-18 (dark red). n ≥ 2. **(H)** Dose-response curves for SN-38 and TAL (CTG, 72 h) in ES8 (black), ES8-KO (red), 2 ES8 BAK KO models (blue), 2 ES8 BAX KO models (green), and 2 ES8 BAK-KAX double KO models (purple). n ≥ 2. **(I)** Heatmap of AUC values in ES8-KO and the BAK, BAX, and BAK-BAX KO models normalized to ES8 (CTG assay, 72 h drug exposure). n ≥ 2. **(J)** Dose-response curves for a 1:1 mixture of SN-38 and TAL combined with either vehicle (DMSO, black), 1 μM venetoclax (“BCL2i”, green), 1 μM S63845 (“MCL1i”, blue), or 1 μM A-1331852 (“BCL-Xli”, red) (CTG, 72 h). n ≥ 2.

Although the *SLFN11* mutation in EW-18 could explain its weak response to DDAs, our analysis of the GDSC and COSMIC databases indicated that the frequency of *SLFN11* mutation in the resistant population was low, with 18/22 cell lines (82%) having a wild-type sequence. We chose SLFN11-wild-type CHLA258 for further investigation. We confirmed strong nuclear expression of the protein by IHC (H-score = 225) (**Fig. 6D**). Consistent with the other tumor lines, SN-38+TAL induced a high level of DNA damage as assessed by the alkaline comet-tail assay (**Fig. 6E**). R-loop expression was comparable to that observed in ES8 and ES8-KO cells (**Fig. 6F**). The “Low” combination of SN-38 and TAL did not substantially increase apoptosis but instead induced a build-up of S-phase cells similar to that observed with ES8-KO cells (**Supplementary Fig. 6SE)**. However, the S-phase population was depleted by the “High” concentration.

Overlaying the SN-38 dose-response curves for the ES8, ES8-KO and resistant cell lines revealed 2 distinct phenotypes (**Fig. 6G**). The dose-response curves for EW-11, EW-13, CHLA258, and A673 (the least sensitive ES cell line in our original panel) showed an inflection at concentrations of <10 nM, as seen with ES8 cells, but did not show the same level of maximum efficacy (percentage inhibition of viability). Consistent with the lack of SLFN11 nuclear localization, EW-18 behaved more like the ES8-KO cells. The efficacy of TAL was substantially less in all cell lines, and its potency was 2 to 6-fold within that in ES8 cells. The exceptions to this were EW-13, in which TAL was more potent than in ES8 cells, and EW-18, in which TAL was inactive (**Supplementary Fig. S6F**).

To probe the mechanism of cytotoxicity induced by TAL and SN-38, we treated ES8 cells exposed to each drug with the caspase inhibitor Z-VAD-FMK, and discovered a significant reduction in efficacy suggesting that apoptosis was the primary mode of cell death with this drug combination (data not shown). To investigate the role of the intrinsic apoptosis pathway in mediating drug sensitivity, we knocked out BAX, BAK, and both BAX and BAK in ES8 cells. The potency of SN-38 and TAL in BAX and BAX/BAK KO cells remained within 2-fold of that in wild-type ES8 cells, but their efficacy was significantly reduced—a phenotype similar to that observed in EW-11, EW-13, CHLA258, and A673 (**Fig. 6H**). BAX deletion alone reduced sensitivity to the DDAs doxorubicin and etoposide to levels comparable to those in ES8-KO cells, but the deletion also reduced sensitivity to vincristine—a phenotype distinct from that of SLFN11 KO cells and comparable to that observed in the resistant ES cell lines described earlier (**Fig. 6I**).

During this project, it was reported that resistance to the PARPi olaparib in ES cell lines could be overcome with the pan-BCL2 inhibitor navitoclax [41]. Motivated by that work and by our own findings, we screened small molecules inhibitors that were selective for individual BCL2 family members: venetoclax (BCL2), S63845 (MCL), and A-1331852 (BCL-XL). BCL-XL inhibition alone sensitized sarcoma cells to the combination of SN-38 and TAL (**Fig. 6J**). Addition of A-1331852 decreased cell viability in A673 and CHLA-258 by a factor of 5.2- and 7.9, respectively, relative to that with the “Low” concentration alone. Although BCL-XL inhibition enhanced the efficacy of the combination in SLFN11-negative U2OS cells, the change in U2OS-OE cells was significantly greater (a 7.5-fold increase vs. a 2.8-fold increase).

These mechanistic studies suggest that (a) impairment of the intrinsic apoptotic pathway, rather than reduced SLFN11 expression, constitutes a primary means of resistance to TAL and SN-38 in pediatric sarcomas; and (b) selective inhibition of BCL-XL can restore sensitivity to the drug combination in SLFN11 expressing resistant tumors.

## DISCUSSION

The heterogeneous and aggressive nature of pediatric sarcomas makes it imperative to identify biomarkers for drug response and new therapeutic targets. Preliminary results from the phase I clinical trial NCT02392793 showed that the combination of TAL and IRN was tolerable and yielded encouraging results in several patients with sarcoma. In a phase I/II trial, TAL+TMZ was also well tolerated; however, no tumor response was seen in the 10 patients with ES treated on that trial, although 2 had prolonged SD [42]. Here, we have shown that SLFN11, not *EWSR1*-fusion or functional BRCA deficiency, is a significant driver of sensitivity to TAL and IRN in this population.

Using IHC, we found that SLFN11 was widely expressed in our cohort of patients with pediatric sarcomas, with H-scores reaching near maximum in some samples. The sensitivity to TAL and SN-38 in sarcoma cell lines correlated with protein levels both *in vitro* and *in vivo* and was independent of the *EWSR1* translocation. However, despite the strong link between protein expression and DDA sensitivity in our preclinical studies, we found no association between SLFN11 status and improved outcome in a retrospective analysis of our patient cohort. Moreover, we found no evidence that SLFN11 was silenced in recurrent disease. These findings contradict recent reports indicating that certain SLFN11-positive tumors have better outcomes with DDA therapy and that gene silencing is a principal route to DDA resistance.

The contrast between our findings and studies in other tumor backgrounds could be attributed to the unique role that SLFN11 plays in pediatric sarcomas. SLFN11 KO in sarcoma cells induced significant changes in transcription, including upregulation of interferon signaling and MHC Class I antigen presentation, and downregulation of genes required for cellular proliferation and oncogenic metabolism. SLFN11-wild-type cells were more sensitive than SLFN11 KO cells to arginine depletion using ADI-PEG-20, confirming the functional consequence of our finding that SLFN11 deletion increased levels of ASS1, the rate-limiting factor in arginine biosynthesis. Moreover, SLFN11 expressing cells grew faster in cell culture, and JN-DSRCT cells engrafted better in orthotopic xenograft models compared to JN-DSCRT-KO cells.

Importantly, we identified sarcoma models expressing high levels of SLFN11 that were resistant to TAL and SN-38. With the exception of EW-18, in which the *SLFN11* gene was truncated, these tumor cell lines expressed wild-type protein as determined by Western blot analysis, sequence analysis, and nuclear localization. We saw no changes in R-loop levels or in the extent of drug-induced DNA damage. These resistant models compensate for the DDA vulnerability induced by SLFN11 expression by attenuating intrinsic apoptosis. Selective inhibition of BCL-XL increased the cytotoxicity of the combination of TAL and SN-38 in resistant ES cell lines, and resulted in enhanced drug efficacy in OST U2OS cells engineered to over-express SLFN11.

Our retrospective analysis of SLFN11 and outcome in pediatric sarcoma was clearly limited. Although most patients received at least 1 DDA, the treatment modalities and disease types varied widely. Moreover, our population appeared to be more refractory to therapy than what would be expected, likely owing to the categories of patients treated at our institution. Consequently, our work strongly supports the prospective evaluation of SLFN11 as a biomarker predicting the response to PARPi and Topo1i combinations in newly diagnosed pediatric sarcomas, which are less likely to have developed widespread chemo-resistance. The ability to rescue tumors by selective BCL-XL inhibition warrants further investigation of strategies that target both replicative stress and the intrinsic apoptotic pathway. Finally, enhanced ASS1 deficiency in SLFN11-positive sarcomas may render them more susceptible to arginine depletion strategies. Ultimately, our work provides a framework to develop rational, targeted combination therapy approaches for both treatment naïve and refractory SLFN11-positive pediatric sarcoma.

## METHODS

### Genomics of Drug Sensitivity in Cancer (GDSC) Correlations

Drug sensitivity (v17.3) and expression data were downloaded from the GDSC website (https://www.cancerrxgene.org/gdsc1000/GDSC1000_WebResources/Home.html) in May 2018 and June 2018, respectively. COSMIC mutation data (CosmicMutantExport.tsv.gz) was downloaded from https://cancer.sanger.ac.uk/cosmic/download in March 2020.

### Mutational Signatures

The set of 30 mutational signatures, a 96 × 30 matrix *Z*, were obtained from COSMIC (https://cancer.sanger.ac.uk/cosmic/signatures). Details of the fitting procedure are provided in Supplemental Data.

### Immunohistochemical staining of pediatric tumor samples

Immunohistochemistry was performed with the Dako Omnis instrument (Agilent) on 4-μm-thick formalin-fixed paraffin-embedded whole-tissue sections, using a rabbit anti-SLFN11 (anti-Schlafen family member 11) polyclonal antibody (Sigma-Aldrich Cat# HPA023030, RRID:AB_1856613) (1:25 dilution, 60 min incubation), Dako Low pH Target Retrieval Solution, and the Dako EnVision Flex Detection Kit. Immunoreactivity was scored using H-scores as described in Supplemental Data.

### Cell culture and viability assays

Description, sourcing, and culture conditions for the sarcoma cell lines used in this work are reported in **Supplementary Table S2A**. Cell lines were authenticated using short tandem repeat analysis via PowerPlex (Promega), and tested for mycoplasma using MycoAlert (Lonza). Translocation status was confirmed using PCR and Fluorescence In Situ Hybridization (FISH). Cell viability was assessed using the CellTiter-Glo (CTG) assay (Promega). ADI-PED 20 was provided by Polaris Pharmaceuticals (San Diego, CA). Cell line characterization by Western Blot and qPCR analysis is described in Supplemental Data.

### Cell Engineering

SLFN11 cDNA (OriGene Technologies Cat# RC226247L4) and the pVector control vector (OriGene Technologies Cat# PS100093) were used to generate the ES8-KO+OE, U2OS-OE, and U2OS-PV models, respectively. hSLFN11^−/−^ cells were generated using CRISPR-Cas9 technology.

### Comet Assay

Alkaline single cell electrophoresis was performed using the CometAssay Reagent kit (Trevigen Cat# 4250-050) in accordance with the manufacturer’s instructions. Comets were imaged by using the LionHeart FX automated microscope (Biotek) and the Gen5 Image Prime software to construct image montages that were analyzed using TriTek CometScore 2.0.0.38.

### R-Loop Immunofluorescence

R-loops were quantified based on the total nuclear intensity of the S9.6 antibody (Kerafast Cat# ENH001, RRID:AB_2687463). Cells were imaged with a Leica microscope using 40x and 63x objectives. Images were acquired using the Photon Counting 3D Nyquist technique.

### Expression Analysis by Microarray

Expression profiles were generated from biological triplicates of wild-type and SLFN11-null cells from two parental cell lines (ES8, JNDS). Cells were treated with 0 or 2 Gy of gamma irradiation then harvested at 4 h or 24 h post-irradiation. Total RNA (100 ng) was purified from treated cells with a RNeasy Mini Kit (Qiagen Cat# 74104) and analyzed using the Affymetrix Clariom S Human assay (ThermoFisher Scientific Cat# 902927).

### Patient Correlations

The St. Jude electronic database was surveyed for solid tumor patients enrolled on St. Jude trials from 2000 to 2018 to identify potential samples. Samples were assessed for viability and availability. Once staining was performed, a retrospective review of the electronic medical record was conducted. Patient data were matched with the IHC samples, and the results were analyzed for correlation by an independent statistician.

### *In Vivo* Experiments

Athymic nude immunodeficient mice were purchased from Charles River (strain code 553). This study was carried out in strict accordance with the recommendations in the Guide to Care and Use of Laboratory Animals of the National Institute of Health. The protocol was approved by the Institutional Animal Care and Use Committee at St. Jude Children’s Research Hospital. Drug dosing and efficacy studies were performed as described previously [6].

## Acknowledgements

The authors thank Keith A. Laycock, PhD, ELS, for scientific editing of the manuscript. We thank Dr. Richard Ashmun and the Flow Cytometry and Cell Sorting Shared Resource at St. Jude for expert guidance, and the Childhood Solid Tumor Network (http://www.stjude.org/CSTN/) for providing xenograft models. Preclinical imaging was performed with help from the Center for In Vivo Therapeutics at St. Jude. Cell images were acquired at the Cell and Tissue Imaging Center, cytogenetics were performed by the Cytogenetic Shared Resource, and microarray data were generated at the Hartwell Center for Bioinformatics and Biotechnology, all of which are supported by St. Jude and by the National Cancer Institute grant P30 CA021765. Additionally, this work was supported by the American Lebanese Syrian Associated Charities (AS, E.A.S.). E.A.S. was supported by the National Comprehensive Cancer Network and is a St. Baldrick’s Scholar with generous support from the Invictus Fund. A.A.S. was supported by the Sarcoma Foundation of America and the St. Baldrick’s Foundation. The content of this paper is solely the responsibility of the authors and does not necessarily represent the official views of the National Institutes of Health.

