## Supplemental Information for "Interrogation of SLFN11 in pediatric sarcomas uncovers an unexpected biological role and a novel therapeutic approach to overcoming resistance to replicative stress"

[1]Department of Oncology, St. Jude Children's Research Hospital, Memphis, Tennessee. [2] Department of Chemical Biology and Therapeutics, St. Jude Children's Research Hospital, Memphis, Tennessee. [3]Department of Pathology, University of Colorado Anschutz Medical Campus, Aurora, Colorado. [4]Department of Pathology, St. Jude Children's Research Hospital, Memphis, Tennessee. [5]Department of Biostatistics, St. Jude Children's Research Hospital, Memphis, Tennessee. [6]Department of Computational Biology, St. Jude Children's Research Hospital, Memphis, Tennessee. [7]Department of Cell and Molecular Biology and The Center for Advanced Genomic Engineering, St. Jude Children's Research Hospital, Memphis, Tennessee. [8]Hartwell Center for Bioinformatics and Biotechnology, St. Jude Children's Research Hospital, Memphis, Tennessee. [9]Department of Radiation Oncology, St. Jude Children's Research Hospital, Memphis, Tennessee. [10]Department of Developmental Neurobiology, St. Jude Children's Research Hospital, Memphis, Tennessee.

Running Title: The role of SLFN11 in pediatric sarcomas

Keywords: SLFN11, pediatrics, sarcoma, replicative stress

Financial support: This work was supported, in part, by the American Lebanese Syrian Associated Charities (E.A.S., A.A.S.). E.A.S. was supported by the National Comprehensive Cancer Network and is a St. Baldrick's Scholar with generous support from the Invictus Fund. A.A.S. was supported by research grants from the Sarcoma Foundation of America and the St. Baldrick's Foundation.

Corresponding authors: (1) Dr. Elizabeth A. Stewart, Department of Oncology and Department of Developmental Neurobiology, St. Jude Children's Research Hospital, 262 Danny Thomas Place, Memphis, TN 38105. Tel: +1 901-595-3544; Fax: +1 901-521-9005;. (2) Dr. Anang A. Shelat, Department of Chemical Biology and Therapeutics, St. Jude Children's Research Hospital, 262 Danny Thomas Place, Memphis, TN 38105. Tel: +1 901-595-5751; Fax: +1 901-595-5715;.

The authors declare there are no conflicts of interest.

Word count = 5928; Total figures and tables = 7

**Table of Contents** **Page**

| | |
|---|---|
| Detailed methods: GDSC correlations | 3 |
| Detailed methods: Mutational signatures | 3 |
| Detailed methods: Procedure for calculating IHC H-score | 4 |
| Detailed methods: Cell viability assay | 4 |
| Detailed methods: Western blot analysis | 6 |
| Detailed methods: Real-time quantitative polymerase chain reaction (RT-qPCR) | 7 |
| Detailed methods: Generation of over-expression models | 7 |
| Detailed methods: Generation of CRISPR knock-out cell lines | 7 |
| Detailed methods: Annexin-V and cell cycle analysis | 9 |
| Detailed methods: Comet assay procedures | 9 |
| Detailed methods: R-loop immunofluorescence | 10 |
| Detailed methods: Expression analysis | 11 |
| Detailed methods: Patient correlation cohort methods | 12 |
| Detailed methods: In vivo experiments | 12 |
| Detailed methods: Analysis of SLFN11 in SJEWS049193 | 15 |
| Description of supplemental tables | 29 |
| Supplemental Figure S1 | 30 |
| Supplemental Figure S2 | 31 |
| Supplemental Figure S3 | 33 |
| Supplemental Figure S4 | 34 |
| Supplemental Figure S5 | 36 |
| Supplemental Figure S6 | 37 |
| References for Supplemental Data | 38 |

#### EXTENDED METHODS

##### GDSC Correlations

We used the area-over-the-curve (AOC) to quantify drug activity, wherein the curve in question is the fitted proportional survival relative to negative controls. This metric gives intuitive results across a wide range of dose-response behaviors, yielding more robust results than those obtained with metrics such as $EC_{50}$ (which can be poorly defined in low-activity systems) or $IC_{50}$ (which is fully undefined for systems that do not reach 50% killing). Fitting proportional survival is done using standard non-linear least-squares regression, fitting a four-parameter log-logistic equation to logarithmically transformed proportional cell survival. Fitting to the logarithm of cell survival allows for more reliable fitting of drugs that achieve high levels of killing (e.g. 1000-fold or 10000-fold killing relative to vehicle). All fits were performed in the R Statistical Computing Environment (1). However, because the AOC in logarithmic survival space is theoretically unbounded, we transformed the fitted survival curve into linear survival space to estimate the AOC to maintain a bounded range of AOC values.

##### Mutational Signatures

To obtain the weight of each signature in a sample, we started with the list of all simple somatic mutations. For a given sample, each single nucleotide mutation was classified as one of the 96 possible mutation types, resulting in a 96 × 1 probability vector $f$. The vector $p$ representing the weights of the 30 COSMIC signatures, was chosen such that the sum $\sum_{i=1}^{96}(Zp - f)^2$ was minimized. The optimization procedure was implemented in MATLAB using the built-in quadratic programming function quadprog.

**Procedure for Calculating IHC H-Score**

Immunoreactivity was scored using H-scores with the percentage of cells with positive staining being estimated at one of 3 levels of intensity (weak, moderate, or strong). Cells with no staining were given a score of 0+. The resultant H-score was calculated with the following formula [H-score = (1 ×%weak)+(2 ×%moderate)+(3 ×%strong)], with the overall score ranging from 0 (negative) to 300 (100% strong staining).

**Cell Viability Assay**

Cell viability was measured using CellTiter-Glo® (Promega). The luminescent signal was read with an EnVision® Multimode Plate Reader (PerkinElmer). The results of screening experiments were processed and visualized using 2 programs developed in-house: RISE (Robust Investigation of Screening Experiments) and AssayExplorer. Cells were plated in 96-well plates at variable densities 12-24 h in advance of drug to ensure that they continued in the log growth phase for the duration of the experiment. The drug plates containing stocks of compounds and the control plate containing the negative control (DMSO) and the positive control (staurosporine) were generated by the Compound Management Center in the Department of Chemical Biology and Therapeutics at our institution. Dose-response experiments were performed with 10-point, 3-fold dilution (19,683-fold concentration range). Before compound transfer, the assay and compound plates were centrifuged at 201 × *g* (1000 rpm) for 1 min in an Eppendorf 5810 centrifuge equipped with an A-4-62 swing-bucket rotor (Eppendorf AG). Approximately 102 nL/well of compound was transferred to a 100 µL volume in the corresponding wells of the assay plates with a 100SS 96-well pin-tool (V&P Scientific) using a Biomek FX Liquid Handler (Beckman Coulter). The final DMSO concentration in each well was ≤0.2%.

Raw luminescence RLU (relative light unit) values for each compound at each concentration were log2 transformed; normalized to obtain the % activity by using the following equation: % activity = 100 × [(mean[negctr]) − compound) / (mean[negctr]) − mean[posctrl])]; then pooled from replicate experiments before fitting. Here, negctrl and posctrl refer to the negative (DMSO) and positive (20–30 µM staurosporine) controls on each plate. Dose-response curves were fit using the drc (2) package in R (1). Both a 3-parameter model (with y0, the response without drug, set to zero) and a 4-parameter model (with y0 allowed to vary) were fit using the sigmoidal function LL2.4. The hill slope was constrained to be between −10 and 0, and the $EC_{50}$ was constrained to be between $10^{-11}$ and $10^{-4}$ (which roughly equated to the drug concentration range tested in these experiments). For the 3-parameter model, yFin, the maximum response of the dose-response curve, was constrained to be between zero and the maximum of the median activities calculated at each concentration over all pooled measurements. For the 4-parameter model, y0 and yFin were both constrained to be between the minimum and the maximum of the median activities calculated at each concentration over all pooled measurements. The model with the lowest corrected Akaike information criterion (AICc) was selected as the best fit model.

The area under the curve (AUC) was calculated from the fitted curve by using the trapezoid rule in the concentration range $10^{-11}$ to $10^{-4}$ molar. In the event of a failure to fit a sigmoidal dose response curve, the smooth.spline option in R was used to fit a curve that could be used to determine the AUC.

**Western Blot Analysis**

The following antibodies were used: BAK (clone D4E4, Cell Signaling Technology Cat# 12105, RRID:AB_2716685), BAX (Cell Signaling Technology Cat# 2772, RRID:AB_10695870), β-Actin (clone 13E5, Cell Signaling Technology Cat# 4970, RRID:AB_2223172), α-Tubulin (clone DM1A, Novus Cat# NB100-690, RRID:AB_521686), IRDye® 680LT Goat anti-Rabbit IgG antibody (LI-COR Biosciences Cat# 926-68021, RRID:AB_10706309), and IRDye® 800CW Donkey anti-mouse IgG antibody (LI-COR Biosciences Cat# 926-32212, RRID:AB_621847). Cell pellets were flash frozen and stored at −80° C. Cells were lysed using radio-immunoprecipitation assay lysis buffer containing cOmplete Mini Protease Inhibitor (Roche) and 1× phosphatase inhibitor, then sonicated at 50% for 20 s and centrifuged at 4°C for 20 min at 20000 × *g*. The supernatant was collected, and the protein concentration was measured using a bicinchoninic acid assay kit (Life Technologies). Proteins were separated by electrophoresis on 10% NuPage Bis-Tris gels and transferred to PVDF membranes (ThermoFisher Scientific), which was then blocked with Odyssey blocking buffer for 1 h. The membranes were incubated with primary antibody and 0.1% Tween overnight at 4°C. Membranes were then washed with PBS and 0.1% Tween (PBST) and secondary antibodies were added with blocking solution and Tween for 40 min. Membranes were then washed with PBST + 0.02% SDS and imaged using an Odyssey CLx Infrared Imaging System (LI-COR). Protein was quantified with LI-COR Biosciences software.

**Real-Time Quantitative Polymerase Chain Reaction (RT-qPCR) Analysis**

Total mRNA was extracted using the Maxwell RSC simplyRNA Tissue Kit and reverse transcribed using iScript Reverse Transcription Supermix (Bio-Rad). TaqMan primers were used for real-time polymerase chain reaction. β-Actin was used as the internal control.

**Generation of Over-Expression Models**

*SLFN11* cDNA (OriGene Technologies Cat# RC226247L4) and pVector control vector (OriGene Technologies Cat# PS100093) were transiently transfected into U2OS cells at 50%–70% confluence by using TurboFectin 8.0 Transfection Reagent (OriGene Technologies) according to the manufacturer's protocol. Twenty-four hours after transfection, cells were passaged into fresh growth medium containing puromycin (selective medium). A mock transfection well in a 6-well plate was used in parallel as a control. The selective medium was changed every 2–3 days until all cells in the control well were dead. The selective medium was then changed into fresh growth medium. The surviving cells were maintained in culture and collected for further validation.

**Generation of CRISPR Knock-out Cell Lines**

hSLFN11$^{-/-}$ cells were generated using CRISPR-Cas9 technology. Briefly, 400,000 cells were transiently co-transfected with 500 ng of gRNA expression plasmid (cloned into Addgene plasmid #43860), 1 µg of Cas9 expression plasmid (Addgene plasmid #43945), and 200 ng of pMaxGFP via nucleofection (Lonza, 4D-Nucleofector™ X Unit), using solution P3 and program DS150 for ES8 cells, EO-100 for A673-shEF cells, and EH-100 for JN-DSRCT-1 cells in small (20 µl) cuvettes in accordance with the manufacturer's recommended protocol. Cells were single-cell sorted by FACS to enrich for GFP+ (transfected) cells, clonally selected, and verified for the desired targeted modification via targeted deep sequencing. Two clones were identified for each modification and assessed in relevant assays. The sequences for the sgRNA and the relevant primers are listed below.

| Name | Sequence (5′ to 3′) |
|---|---|
| hSLFN11 sgrna | UGUCAGCUGAGUCUAUCUAG |
| hSLFN11.NGS.F<br><br>partial Illumina adaptors (upper case) | CACTCTTTCCCTACACGACGCTCTTCCGATCTaagtcacacattccacctactga |
| hSLFN11.NGS.R<br><br>partial Illumina adaptors (upper case) | GTGACTGGAGTTCAGACGTGTGCTCTTCCGATCTggtttgcagagcactttcagg |

<u>ES8 hSLFN11 KO</u>

Clone 1F2

−1bp: 5′-AAGATTTTGAATGTCAGCTGAGTCTATCAGTGGGCCTCCCCTTAGCAGACC-3′

+1bp: 5′-AAGATTTTGAATGTCAGCTGAGTCTATC**C**TAGTGGGCCTCCCCTTAGCAGACC-3′

<u>A673-shEF hSLFN11 KO</u>

Clone 2B9

−4bp: 5′-AAGATTTTGAATGTCAGCTGAGTCTA----GTGGGCCTCCCCTTAGCAGACC-3′

<u>JN-DSCRT-1-YFP hSLFN11 KO</u>

Clone 3B5

−4bp: 5′-AAGATTTTGAATGTCAGCTGAGTCTA----GTGGGCCTCCCCTTAGCAGACC-3′

−1bp: 5′-AAGATTTTGAATGTCAGCTGAGTCTAT-TAGTGGGCCTCCCCTTAGCAGACC-3′

**Annexin-V and Cell Cycle Analysis**

Cells were plated at appropriate densities to obtain $5x10^5$–$1x10^6$ cells at the time of collection and allowed to adhere 24 h at 37°C prior to addition of drug. After drug treatment, the supernatant was collected and combined with gently trypsinized cells that were then resuspended in fresh medium. Cells were then divided equally and placed on ice for cell cycle analysis and quantification of apoptosis.

Half of the cells were stained with annexin-V and DAPI to determine their apoptosis levels. The other half were stained with propidium iodide to determine their cell cycle status. Cells were interrogated using an LSR Fortessa Flow Cytometer (BD Bioscience). Apoptosis statistics were obtained with DiVa Software (BD Bioscience), and cell cycle statistics were obtained with ModFit software (Verity Software House).

**Comet Assay Procedures**

Alkaline single-cell electrophoresis was performed using the CometAssay Reagent Kit (Trevigen Cat# 4250-050) in accordance with the manufacturer's instructions. Briefly, cells were plated at a density of 50,000 cells per well in a 96-well plate, incubated for 24 h, then treated with 0.2%

DMSO, 1 µM TAL, 1 µM SN-38, or a combination of 1 µM TAL and 1 µM SN-38. After being treated for 2.5 h, the cells were trypsinized then washed with ice-cold 1× PBS. They were then combined with low-melting-point agarose (LMAgarose, Trevigen Cat# 4250-500-02) at a ratio of 1:10 (v/v) and immediately plated onto CometSlides (Trevigen Cat# 4253-096-03). The slides were placed at 4°C in the dark for 30 min to improve cell adherence then immersed in lysis solution for 60 min. The slides were then drained and incubated with Alkaline Unwinding Solution for 20 min at room temperature. Electrophoresis was then performed using the CometAssay® Electrophoresis System II unit (Trevigen Cat# 4250-050-ES) with 21 volts being applied for 30 min for ES8 and ES8-KO cells and for 45 min for CHLA-258. Samples were dried for 30 min at 37°C and washed with $dH_2O$ × 2 and 70% ethanol. Samples were then dried for an additional 30 min at 37°C then stained with SYBR® Gold solution for 30 min.

**R-Loop Immunofluorescence**

Cells were seeded on Ibidi slides (uClear) and allowed to attach overnight. Next day, the cells were fixed using cold, dry methanol for 30 min at 4°C followed by a wash step with cold phosphate-buffered saline (PBS). The cells were then permeabilized with 10% normal goat serum (NGS) and 0.1% Triton-X in PBS for 20 min at room temperature (RT). They were then washed with PBS and incubated with S9.6 primary antibody (Kerafast Cat# ENH001, RRID:AB_2687463) in 1% NGS in PBS overnight. After washing with PBS, the cells were incubated with Alexa Fluor, Goat anti-Mouse, 647 antibody (Molecular Probes) in 1% NGS in PBS at RT for 2 h then stained with Hoechst stain (Invitrogen). Cells were imaged with a Leica microscope using 40× and 63× objectives. Images were collected using the Photon Counting 3D Nyquist technique. To quantify R-loop levels, nuclear outlines were traced by hand, using the freeform selection tool in ImageJ, on images of dimensions 1084 pixels × 1084 pixels, representing regions of 150 µm × 150 µm. To calculate the background-subtracted R9.6 signal, the R9.6 channel of each image was blurred with a Gaussian filter with a standard deviation of 2 pixels (to smooth out dead or outlier pixels) then passed through a local minimum filter with a radius of 20 pixels. This gave the minimum nearby value for each pixel and was subtracted from the original R9.6 channel. The mean image intensity and area were calculated on the background-subtracted R9.6 channel by using the Measurement tool in ImageJ. The total nucleolar luminous intensity was estimated by multiplying the average background-subtracted R9.6 intensity (ranging from 0 to 1) by the area of the nucleus (scaled to square micrometers).

**Expression Analysis**

Probe signals were normalized and transformed into log2 transcript expression values by the robust multi-array average (RMA) algorithm, using the Affymetrix Expression Console software v1.1. Differentially expressed transcripts were identified by an ANOVA model in which parental lineage, genotype, radiation dose, and irradiation timepoint were used as the factors (Partek Genomics Suite software v6.6; Partek, Inc.). The false-discovery rate (FDR) was estimated by the method of Benjamini and Hochberg, with an FDR threshold of <0.05 being applied to identify differentially expressed transcripts. Gene lists were analyzed for pathway and functional enrichment by using Ingenuity Pathways Analysis (Qiagen). Differential expression was also investigated using Gene Set Enrichment Analysis (GSEA). However, in our initial attempts to identify genotype or treatment effects common to both cell lineages, the results were obscured by the magnitude of expression differences between the cell lineages. This dominant effect of cell lineage was also confirmed by principal component analysis and by the ANOVA model. Therefore, to identify genotype or treatment differences common to both lineages, we performed GSEA after adjusting the dataset for the effects of cell lineage using the 4-factor ANOVA model in Partek Genomics Suite software v6.6. To validate the use of the lineage-adjusted data for GSEA, we also performed GSEA using the pre-ranked method, in which gene rankings were determined by the ratio of expression profiles determined by the 4-factor ANOVA model. The GSEA results from both pre-ranked and lineage-adjusted data were virtually identical.

**Patient Correlation Cohort Methods**

Data collection resulted in a dataset of 353 pathology samples from 220 patients. Samples obtained at the same time from the same specimen were summarized by their median value, which reduced the number of sample measurements to 335. We defined a patient as “SLFN11 positive” if any sample from that patient had an H-score > 0 at any time, and “SLFN11 negative” if every sample from that patient had an H-score = 0. We restricted the survival analysis to patients who had a sample at diagnosis or treatment. The observation selected for inclusion in this data subset was the first of the following: (1) human sample at diagnosis, (2) xenograft sample at diagnosis, (3) human sample at treatment, and (4) xenograft sample at treatment. If a patient did not have any observation from the above list, the patient was excluded from the data subset. This data subset contained 143 patients (one measurement per patient).

**In vivo Experiments**

All efforts were made to minimize suffering. All mice were housed in accordance with approved IACUC protocols. Animals were housed on a 12-12 light cycle (lights on at 6am, off at 6pm) and provided with food and water *ad libitum*.

Orthotopic xenografts were created by injecting luciferase-labeled cells into athymic nude mice, using the following techniques: bone marrow injection, as described previously (3), for ES8, ES8-KO, and 143B cells; intraperitoneal injection for JN-DSRCT-1 and JN-DSRCT-1 KO cells; and subcutaneous injection for SU-CCS-1 cells. SJEWS049193_X1 was derived from an 11-year-old male patient with a history of recurrent metastatic Ewing sarcoma under the MAST protocol (NCT01050296). Patient-derived xenografts were created using SJEWS049193_X1 tumor cells obtained from the Childhood Solid Tumor Network (http://www.stjude.org/CSTN/). These cells were dissociated and passaged as described previously (4).

Mice were screened weekly by Xenogen and the bioluminescence was measured. Mice were enrolled in the study after a target bioluminescence signal of $10^7$ photons/s/$cm^2$ or a palpable tumor was obtained, and chemotherapy was started on the following Monday. Treatment schedules and dosing are described below. Mice received 4 courses of chemotherapy (3 weeks per course) and bioluminescence was monitored weekly and at the end of therapy. Mice were monitored daily while receiving chemotherapy. Disease response was classified according to the bioluminescence signal (see table below).

<u>Treatment groups for *in vivo* experiments</u>

| Cell line | No. of treatment groups | Treatment groups |
|---|---|---|
| ES8 | 4 | Control, TAL+IRN, IRN+TMZ, TAL$^{\pm}$+IRN+TMZ |
| ES8-KO | 4 | Control, TAL+IRN, IRN+TMZ, TAL$^{\pm}$+IRN+TMZ |
| JN-DSRCT-1 | 3 | Control, TAL+IRN, TAL$^{\pm}$+IRN+TMZ$^{\pm}$ |
| 143B | 3 | Control, TAL+IRN, TAL$^{\pm}$+IRN+TMZ$^{\pm}$ |
| SU-CCS-1 | 3 | Control, TAL+IRN, TAL$^{\pm}$+IRN+TMZ$^{\pm}$ |

| Drug | Dose and schedule |
|---|---|
| IRN | 1.25 mg/kg by IP injection once daily on days 1–5, 8–12 |
| TAL | 0.125 mg/kg by oral gavage twice daily on days 1–5, 8–12 |
| TAL$^{\pm}$ | 0.1 mg/kg by oral gavage twice daily on days 1–5, 8–12 |
| TMZ | 33 mg/kg by oral gavage once daily on days 1–5 |
| TMZ$^{\pm}$ | 10 mg/kg by oral gavage once daily on days 1–5 |
| Control | No chemotherapy |

$^{\pm}$dose reduced to limit toxicity

<u>Response criteria for *in vivo* experiments</u>

*Xenogen® Imaging and Quantification*

Mice were given intraperitoneal injections of Firefly D-Luciferin (Caliper Life Sciences) (3 mg/mouse). Bioluminescent images were acquired 5 minutes later with the IVIS® 200 imaging system. Anesthesia (isoflurane 1.5% in $O_2$ delivered at 2 L/min) was administered throughout image acquisition.  Living Image 4.3 software (Caliper Life Sciences) was used to generate a standard region of interest (ROI) encompassing the largest tumor at the maximal bioluminescence signal. The identical ROI was used to determine the average radiance (in photons/s/$cm^2$/sr) for all xenografts. Response was defined as below:

| **Response** | **Xenogen signal** |
|---|---|
| Complete response | ≤$10^5$ photons/s/$cm^2$ (similar to background) |
| Partial response | $10^5$–$10^7$ photons/s/$cm^2$ |
| Stable disease | $10^7$–$10^8$ photons/s/$cm^2$ (similar to the enrollment signal) |
| Progressive Disease* | >$10^8$ photons/s/$cm^2$ |

*Mice with tumor burden greater than 20% of body weight at any time were also classified as having progressive disease.

Growth curves for untreated controls

ES8 and ES8-KO: Nude mice were injected intrafemorally with luciferase-labeled cells (1 million cells/mouse). As these tumors are known to engraft rapidly in mice, Xenogen imaging was started within 1–2 weeks of injection and continued weekly.

JN-DSRCT and JN-DSRCT-KO: Nude mice were injected subcutaneously with luciferase-labeled cells (1 million cells/mouse) because previous attempts to engraft JN-DSRCT KO cells in the intraperitoneal space were unsuccessful. As these tumors are slower growing in mice compared to other ES8, Xenogen imaging was started 3–4 weeks after injection and continued weekly.

**Analysis of *SLFN11* in SJEWS049193**

Consensus RNA-seq SLFN11 sequences (chr17:33675329-33702720) from SJEWS049193_D1 (primary site tumor, site #1), SJEWS049193_D2 (primary site tumor, site #2), SJEWS049193_X1 (metastatic tumor, site #1), and SJEWS049193_X2 (metastatic tumor, site #2) were aligned to the coding region (exons 4, 5, 6, and 7) of *hSLFN11* (NM_001104587). The SLFN11 sequences from these 4 independent tumor samples have wild-type sequences but share 4 known SNPs:

1. rs4796077 (Asn301Asp): $^{901}$ AAT (Asn) → GAT (Asp)
2. rs28571667 (Synonymous, Gln479): $^{1435}$ CAA (Gln) → CAG (Gln)
3. rs9898983 (Arg489Leu): $^{1465}$ CGC (Arg) → CTC (Leu)
4. rs9897352 (Synonymous, Thr490): $^{1468}$ ACC (Thr) → ACT (Thr)

### SJEWS049193_X2

#### Exon 4

**Sequence ID: Query_51559  Length: 27392  Number of Matches: 2**

**Range 1: 14430 to 15498** Graphics    ▼ Next Match  ▲ Previous Match

| Score | Expect | Identities | Gaps | Strand |
|---|---|---|---|---|
| 1924 bits(2133) | 0.0 | 1068/1069(99%) | 0/1069(0%) | Plus/Minus |

```
Query  1      ATGGAGGCAAATCAGTGCCCCCTGGTTGTGGAACCATCTTACCCAGACCTGGTCATCAAT  60
Sbjct  15498  ............................................................  15439

Query  61     GTAGGAGAAGTGACTCTTGgagaagaaaacagaaaaaagctgcagaaaattcagagagac  120
Sbjct  15438  ............................................................  15379

Query  121    caagagaaggagagagTTATGCGGGCTGCATGTGCTTTATTAAACTCAGGAGGAGGAGTG  180
Sbjct  15378  ............................................................  15319

Query  181    ATTCGAATGGCCAAGAAGGTTGAGCATCCCGTGGAGATGGGACTGGATTTAGAACAGTCT  240
Sbjct  15318  ............................................................  15259

Query  241    TTGAGAGAGCTTATTCAGTCTTCAGATCTGCAGGCTTTCTTTGAGACCAAGCAACAAGGA  300
Sbjct  15258  ............................................................  15199

Query  301    AGGTGTTTTTACATTTTTGTTAAATCTTGGAGCAGTGGCCCTTTCCCTGAAGATCGCTCT  360
Sbjct  15198  ............................................................  15139

Query  361    GTCAAGCCCCGCCTTTGCAGCCTCAGTTCTTCATTATACCGTAGATCTGAGACCTCTGTG  420
Sbjct  15138  ............................................................  15079

Query  421    CGTTCCATGGACTCAAGAGAGGCATTCTGTTTCCTGAAGACCAAAAGGAAGCCAAAAATC  480
Sbjct  15078  ............................................................  15019

Query  481    TTGGAAGAAGGACCTTTTCACAAAATTCACAAGGGTGTATACCAAGAGCTCCCTAACTCG  540
Sbjct  15018  ............................................................  14959

Query  541    GATCCTGCTGACCCAAACTCGGATCCTGCTGACCTAATTTTCCAAAAAGACTATCTTGAA  600
Sbjct  14958  ............................................................  14899

Query  601    TATGGTGAAATCCTGCCTTTTCCTGAGTCTCAGTTAGTAGAGTTTAAACAGTTCTCTACA  660
Sbjct  14898  ............................................................  14839

Query  661    AAACACTTCCAAGAATATGTAAAAAGGACAATTCCAGAATACGTCCCTGCATTTGCAAAC  720
Sbjct  14838  ............................................................  14779

Query  721    ACTGGAGGAGGCTATCTTTTTATTGGAGTGGATGATAAGAGTAGGGAAGTCCTGGGATGT  780
Sbjct  14778  ............................................................  14719

Query  781    GCAAAAGAAAATGTTGACCCTGACTCTTTGAGAAGGAAAATAGAACAAGCCATATACAAA  840
Sbjct  14718  ............................................................  14659

Query  841    CTACCTTGTGTTCATTTTTGCCAACCCCAACGCCCGATAACCTTCACACTCAAAATTGTG  900
Sbjct  14658  ............................................................  14599

Query  901    AATGTGTTAAAAAGGGGAGAGCTCTATGGCTATGCTTGCATGATCAGAGTAAATCCCTTC  960
Sbjct  14598  G...........................................................  14539

Query  961    TGCTGTGCAGTGTTCTCAGAAGCTCCCAATTCATGGATAGTGGAGGACAAGTACGTCTGC  1020
Sbjct  14538  ............................................................  14479

Query  1021   AGCCTGACAACCGAGAAATGGGTAGGCATGATGACAGACACAGATCCAG  1069
Sbjct  14478  .................................................  14430
```

#### Exon 5

**Sequence ID: Query_45909** **Length: 27392** **Number of Matches: 1**

**Range 1: 11934 to 12062** Graphics ▼ Next Match ▲ Previous Match

| Score | Expect | Identities | Gaps | Strand |
|---|---|---|---|---|
| 233 bits(258) | 1e-64 | 129/129(100%) | 0/129(0%) | Plus/Minus |

```
Query  1      ATCTTCTACAGTTGTCTGAAGATTTTGAATGTCAGCTGAGTCTATCTAGTGGGCCTCCCC  60
Sbjct  12062  ............................................................  12003

Query  61     TTAGCAGACCAGTGTACTCCAAGAAAGGCCTGGAACATAAAAAGGAACTCCAGCAACTTT  120
Sbjct  12002  ............................................................  11943

Query  121    TATTTTCAG  129
Sbjct  11942  .........  11934
```

#### Exon 6

**Sequence ID: Query_25601** **Length: 27392** **Number of Matches: 1**

**Range 1: 5027 to 5750** Graphics ▼ Next Match ▲ Previous Match

| Score | Expect | Identities | Gaps | Strand |
|---|---|---|---|---|
| 1293 bits(1433) | 0.0 | 721/724(99%) | 0/724(0%) | Plus/Minus |

```
Query  1     TCCCACCAGGATATTTGCGATATACTCCAGAGTCACTCTGGAGGGACCTGATCTCAGAGC  60
Sbjct  5750  ............................................................  5691

Query  61    ACAGAGGACTAGAGGAGTTAATAAATAAGCAAATGCAACCTTTCTTTCGGGGAATTTTGA  120
Sbjct  5690  ............................................................  5631

Query  121   TCTTCTCTAGAAGTTGGGCTGTGGACCTGAACTTGCAGGAGAAGCCAGGAGTCATCTGTG  180
Sbjct  5630  ............................................................  5571

Query  181   ATGCTCTGCTGATAGCACAGAACAGCACCCCCATTCTCTACACCATTCTCAGGGAGCAAG  240
Sbjct  5570  ..........................................................G.  5511

Query  241   ATGCAGAGGGCCAGGACTACTGCACTCGCACCGCCTTTACTTTGAAGCAGAAGCTAGTGA  300
Sbjct  5510  ...........................T...T............................  5451

Query  301   ACATGGGGGGCTACACCGGGAAGGTGTGTGTCAGGGCCAAGGTCCTCTGCCTGAGTCCTG  360
Sbjct  5450  ............................................................  5391

Query  361   AGAGCAGCGCAGAGGCCTTGGAGGCTGCAGTGTCTCCGATGGATTACCCTGCGTCCTATA  420
Sbjct  5390  ............................................................  5331

Query  421   GCCTTGCAGGCACCCAGCACATGGAAGCCCTGCTGCAGTCCCTCGTGATTGTCTTACTCG  480
Sbjct  5330  ............................................................  5271

Query  481   GCTTCAGGTCTCTCTTGAGTGACCAGCTCGGCTGTGAGGTTTTAAATCTGCTCACAGCCC  540
Sbjct  5270  ............................................................  5211

Query  541   AGCAGTATGAGATATTCTCCAGAAGCCTCCGCAAGAACAGAGAGTTGTTTGTCCACGGCT  600
Sbjct  5210  ............................................................  5151

Query  601   TACCTGGCTCAGGGAAGACCATCATGGCCATGAAGATCATGGAGAAGATCAGGAATGTGT  660
Sbjct  5150  ............................................................  5091

Query  661   TTCACTGTGAGGCACACAGAATTCTCTACGTTTGTGAAAACCAGCCTCTGAGGAACTTTA  720
Sbjct  5090  ............................................................  5031

Query  721   TCAG  724
Sbjct  5030  ....  5027
```

#### Exon 7

**Sequence ID: Query_47887 Length: 27392 Number of Matches: 1**

**Range 1: 4047 to 4830** Graphics ▼ Next Match ▲ Previous Match

| Score | Expect | Identities | Gaps | Strand |
|---|---|---|---|---|
| 1415 bits(1568) | 0.0 | 784/784(100%) | 0/784(0%) | Plus/Minus |

```
Query  1     TGATAGAAATATCTGCCGAGCAGAGACCCGGAAAACTTTCCTAAGAGAAAACTTTGAACA  60
Sbjct  4830  ............................................................  4771

Query  61    CATTCAACACATCGTCATTGACGAAGCTCAGAATTTCCGTACTGAAGATGGGGACTGGTA  120
Sbjct  4770  ............................................................  4711

Query  121   TGGGAAGGCAAAAAGCATCACTCGGAGAGCAAAGGGTGGCCCAGGAATTCTCTGGATCTT  180
Sbjct  4710  ............................................................  4651

Query  181   TCTGGATTACTTTCAGACCAGCCACTTGGATTGCAGTGGCCTCCCTCCTCTCTCAGACCA  240
Sbjct  4650  ............................................................  4591

Query  241   ATATCCAAGAGAAGAGCTCACCAGAATAGTTCGCAATGCAGATCCAATAGCCAAGTACTT  300
Sbjct  4590  ............................................................  4531

Query  301   ACAAAAAGAAATGCAAGTAATTAGAAGTAATCCTTCATTTAACATCCCCACTGGGTGCCT  360
Sbjct  4530  ............................................................  4471

Query  361   CGAGGTATTTCCTGAAGCCGAATGGTCCCAGGGTGTTCAGGGAACCTTACGAATTAAGAA  420
Sbjct  4470  ............................................................  4411

Query  421   ATACTTGACTGTGGAGCAAATAATGACCTGTGTGGCAGACACGTGCAGGCGCTTCTTTGA  480
Sbjct  4410  ............................................................  4351

Query  481   TAGGGGCTATTCTCCAAAGGATGTTGCTGTGCTTGTCAGCACCGCAAAAGAAGTGGAGCA  540
Sbjct  4350  ............................................................  4291

Query  541   CTATAAGTATGAGCTCTTGAAAGCAATGAGGAAGAAAAGGGTGGTGCAGCTCAGTGATGC  600
Sbjct  4290  ............................................................  4231

Query  601   ATGTGATATGTTGGGTGATCACATTGTGTTGGACAGTGTTCGGCGATTCTCAGGCCTGGA  660
Sbjct  4230  ............................................................  4171

Query  661   AAGGAGCATAGTGTTTGGGATCCATCCAAGGACAGCTGACCCAGCTATCTTACCCAATGT  720
Sbjct  4170  ............................................................  4111

Query  721   TCTGATCTGTCTGGCTTCCAGGGCAAAACAACACCTGTATATTTTTCCGTGGGGTGGCCA  780
Sbjct  4110  ............................................................  4051

Query  781   TTAG  784
Sbjct  4050  ....  4047
```

### SJEWS049193_X1

#### Exon 4

Sequence ID: **Query_43683** Length: **27392** Number of Matches: **2**

**Range 1: 14430 to 15498** Graphics ▼ Next Match ▲ Previous M

| Score | Expect | Identities | Gaps | Strand |
|---|---|---|---|---|
| 1924 bits(2133) | 0.0 | 1068/1069(99%) | 0/1069(0%) | Plus/Minus |

```
Query  1      ATGGAGGCAAATCAGTGCCCCCTGGTTGTGGAACCATCTTACCCAGACCTGGTCATCAAT  60
Sbjct  15498  ............................................................  15439

Query  61     GTAGGAGAAGTGACTCTTGgagaagaaaacagaaaaaagctgcagaaaattcagagagac  120
Sbjct  15438  ............................................................  15379

Query  121    caagagaaggagagagTTATGCGGGCTGCATGTGCTTTATTAAACTCAGGAGGAGGAGTG  180
Sbjct  15378  ............................................................  15319

Query  181    ATTCGAATGGCCAAGAAGGTTGAGCATCCCGTGGAGATGGGACTGGATTTAGAACAGTCT  240
Sbjct  15318  ............................................................  15259

Query  241    TTGAGAGAGCTTATTCAGTCTTCAGATCTGCAGGCTTTCTTTGAGACCAAGCAACAAGGA  300
Sbjct  15258  ............................................................  15199

Query  301    AGGTGTTTTTACATTTTTGTTAAATCTTGGAGCAGTGGCCCTTTCCCTGAAGATCGCTCT  360
Sbjct  15198  ............................................................  15139

Query  361    GTCAAGCCCCGCCTTTGCAGCCTCAGTTCTTCATTATACCGTAGATCTGAGACCTCTGTG  420
Sbjct  15138  ............................................................  15079

Query  421    CGTTCCATGGACTCAAGAGAGGCATTCTGTTTCCTGAAGACCAAAAGGAAGCCAAAAATC  480
Sbjct  15078  ............................................................  15019

Query  481    TTGGAAGAAGGACCTTTTCACAAAATTCACAAGGGTGTATACCAAGAGCTCCCTAACTCG  540
Sbjct  15018  ............................................................  14959

Query  541    GATCCTGCTGACCCAAACTCGGATCCTGCTGACCTAATTTTCCAAAAAGACTATCTTGAA  600
Sbjct  14958  ............................................................  14899

Query  601    TATGGTGAAATCCTGCCTTTTCCTGAGTCTCAGTTAGTAGAGTTTAAACAGTTCTCTACA  660
Sbjct  14898  ............................................................  14839

Query  661    AAACACTTCCAAGAATATGTAAAAAGGACAATTCCAGAATACGTCCCTGCATTTGCAAAC  720
Sbjct  14838  ............................................................  14779

Query  721    ACTGGAGGAGGCTATCTTTTTATTGGAGTGGATGATAAGAGTAGGGAAGTCCTGGGATGT  780
Sbjct  14778  ............................................................  14719

Query  781    GCAAAAGAAAATGTTGACCCTGACTCTTTGAGAAGGAAAATAGAACAAGCCATATACAAA  840
Sbjct  14718  ............................................................  14659

Query  841    CTACCTTGTGTTCATTTTTGCCAACCCCAACGCCCGATAACCTTCACACTCAAAATTGTG  900
Sbjct  14658  ............................................................  14599

Query  901    AATGTGTTAAAAAGGGGAGAGCTCTATGGCTATGCTTGCATGATCAGAGTAAATCCCTTC  960
Sbjct  14598  G...........................................................  14539

Query  961    TGCTGTGCAGTGTTCTCAGAAGCTCCCAATTCATGGATAGTGGAGGACAAGTACGTCTGC  1020
Sbjct  14538  ............................................................  14479

Query  1021   AGCCTGACAACCGAGAAATGGGTAGGCATGATGACAGACACAGATCCAG  1069
Sbjct  14478  .................................................  14430
```

#### Exon 5

**Sequence ID: Query_15111 Length: 27392 Number of Matches: 1**

**Range 1: 11934 to 12062** Graphics ▼ Next Match ▲ Previous

| Score | Expect | Identities | Gaps | Strand |
|---|---|---|---|---|
| 233 bits(258) | 1e-64 | 129/129(100%) | 0/129(0%) | Plus/Minus |

```
Query  1      ATCTTCTACAGTTGTCTGAAGATTTTGAATGTCAGCTGAGTCTATCTAGTGGGCCTCCCC  60
Sbjct  12062  ............................................................  12003

Query  61     TTAGCAGACCAGTGTACTCCAAGAAAGGCCTGGAACATAAAAAGGAACTCCAGCAACTTT  120
Sbjct  12002  ............................................................  11943

Query  121    TATTTTCAG  129
Sbjct  11942  .........  11934
```

#### Exon 6

**Sequence ID: Query_44151 Length: 27392 Number of Matches: 1**

**Range 1: 5027 to 5750** Graphics ▼ Next Match ▲ Previous M

| Score | Expect | Identities | Gaps | Strand |
|---|---|---|---|---|
| 1293 bits(1433) | 0.0 | 721/724(99%) | 0/724(0%) | Plus/Minus |

```
Query  1     TCCCACCAGGATATTTGCGATATACTCCAGAGTCACTCTGGAGGGACCTGATCTCAGAGC  60
Sbjct  5750  ............................................................  5691

Query  61    ACAGAGGACTAGAGGAGTTAATAAATAAGCAAATGCAACCTTTCTTTCGGGGAATTTTGA  120
Sbjct  5690  ............................................................  5631

Query  121   TCTTCTCTAGAAGTTGGGCTGTGGACCTGAACTTGCAGGAGAAGCCAGGAGTCATCTGTG  180
Sbjct  5630  ............................................................  5571

Query  181   ATGCTCTGCTGATAGCACAGAACAGCACCCCCATTCTCTACACCATTCTCAGGGAGCAAG  240
Sbjct  5570  ..........................................................G.  5511

Query  241   ATGCAGAGGGCCAGGACTACTGCACTCGCACCGCCTTTACTTTGAAGCAGAAGCTAGTGA  300
Sbjct  5510  ...........................T...T............................  5451

Query  301   ACATGGGGGGCTACACCGGGAAGGTGTGTGTCAGGGCCAAGGTCCTCTGCCTGAGTCCTG  360
Sbjct  5450  ............................................................  5391

Query  361   AGAGCAGCGCAGAGGCCTTGGAGGCTGCAGTGTCTCCGATGGATTACCCTGCGTCCTATA  420
Sbjct  5390  ............................................................  5331

Query  421   GCCTTGCAGGCACCCAGCACATGGAAGCCCTGCTGCAGTCCCTCGTGATTGTCTTACTCG  480
Sbjct  5330  ............................................................  5271

Query  481   GCTTCAGGTCTCTCTTGAGTGACCAGCTCGGCTGTGAGGTTTTAAATCTGCTCACAGCCC  540
Sbjct  5270  ............................................................  5211

Query  541   AGCAGTATGAGATATTCTCCAGAAGCCTCCGCAAGAACAGAGAGTTGTTTGTCCACGGCT  600
Sbjct  5210  ............................................................  5151

Query  601   TACCTGGCTCAGGGAAGACCATCATGGCCATGAAGATCATGGAGAAGATCAGGAATGTGT  660
Sbjct  5150  ............................................................  5091

Query  661   TTCACTGTGAGGCACACAGAATTCTCTACGTTTGTGAAAACCAGCCTCTGAGGAACTTTA  720
Sbjct  5090  ............................................................  5031

Query  721   TCAG  724
Sbjct  5030  ....  5027
```

#### Exon 7

**Sequence ID: Query_14007 Length: 27392 Number of Matches: 1**

**Range 1: 4047 to 4830** Graphics ▼ Next Match ▲ Previous Match

| Score | Expect | Identities | Gaps | Strand |
|---|---|---|---|---|
| 1415 bits(1568) | 0.0 | 784/784(100%) | 0/784(0%) | Plus/Minus |

```
Query  1     TGATAGAAATATCTGCCGAGCAGAGACCCGGAAAACTTTCCTAAGAGAAAACTTTGAACA  60
Sbjct  4830  ............................................................  4771

Query  61    CATTCAACACATCGTCATTGACGAAGCTCAGAATTTCCGTACTGAAGATGGGGACTGGTA  120
Sbjct  4770  ............................................................  4711

Query  121   TGGGAAGGCAAAAAGCATCACTCGGAGAGCAAAGGGTGGCCCAGGAATTCTCTGGATCTT  180
Sbjct  4710  ............................................................  4651

Query  181   TCTGGATTACTTTCAGACCAGCCACTTGGATTGCAGTGGCCTCCCTCCTCTCTCAGACCA  240
Sbjct  4650  ............................................................  4591

Query  241   ATATCCAAGAGAAGAGCTCACCAGAATAGTTCGCAATGCAGATCCAATAGCCAAGTACTT  300
Sbjct  4590  ............................................................  4531

Query  301   ACAAAAAGAAATGCAAGTAATTAGAAGTAATCCTTCATTTAACATCCCCACTGGGTGCCT  360
Sbjct  4530  ............................................................  4471

Query  361   CGAGGTATTTCCTGAAGCCGAATGGTCCCAGGGTGTTCAGGGAACCTTACGAATTAAGAA  420
Sbjct  4470  ............................................................  4411

Query  421   ATACTTGACTGTGGAGCAAATAATGACCTGTGTGGCAGACACGTGCAGGCGCTTCTTTGA  480
Sbjct  4410  ............................................................  4351

Query  481   TAGGGGCTATTCTCCAAAGGATGTTGCTGTGCTTGTCAGCACCGCAAAAGAAGTGGAGCA  540
Sbjct  4350  ............................................................  4291

Query  541   CTATAAGTATGAGCTCTTGAAAGCAATGAGGAAGAAAAGGGTGGTGCAGCTCAGTGATGC  600
Sbjct  4290  ............................................................  4231

Query  601   ATGTGATATGTTGGGTGATCACATTGTGTTGGACAGTGTTCGGCGATTCTCAGGCCTGGA  660
Sbjct  4230  ............................................................  4171

Query  661   AAGGAGCATAGTGTTTGGGATCCATCCAAGGACAGCTGACCCAGCTATCTTACCCAATGT  720
Sbjct  4170  ............................................................  4111

Query  721   TCTGATCTGTCTGGCTTCCAGGGCAAAACAACACCTGTATATTTTTCCGTGGGGTGGCCA  780
Sbjct  4110  ............................................................  4051

Query  781   TTAG  784
Sbjct  4050  ....  4047
```

### SJEWS049193_D2

#### Exon 4

Sequence ID: **Query_54525** Length: **27392** Number of Matches: **2**

**Range 1: 14430 to 15498** Graphics ▼ Next Match ▲ Previ

| Score | Expect | Identities | Gaps | Strand |
|---|---|---|---|---|
| 1924 bits(2133) | 0.0 | 1068/1069(99%) | 0/1069(0%) | Plus/Minus |

```
Query  1      ATGGAGGCAAATCAGTGCCCCCTGGTTGTGGAACCATCTTACCCAGACCTGGTCATCAAT  60
Sbjct  15498  ............................................................  15439

Query  61     GTAGGAGAAGTGACTCTTGgagaagaaaacagaaaaaagctgcagaaaattcagagagac  120
Sbjct  15438  ............................................................  15379

Query  121    caagagaaggagagagTTATGCGGGCTGCATGTGCTTTATTAAACTCAGGAGGAGGAGTG  180
Sbjct  15378  ............................................................  15319

Query  181    ATTCGAATGGCCAAGAAGGTTGAGCATCCCGTGGAGATGGGACTGGATTTAGAACAGTCT  240
Sbjct  15318  ............................................................  15259

Query  241    TTGAGAGAGCTTATTCAGTCTTCAGATCTGCAGGCTTTCTTTGAGACCAAGCAACAAGGA  300
Sbjct  15258  ............................................................  15199

Query  301    AGGTGTTTTTACATTTTTGTTAAATCTTGGAGCAGTGGCCCTTTCCCTGAAGATCGCTCT  360
Sbjct  15198  ............................................................  15139

Query  361    GTCAAGCCCCGCCTTTGCAGCCTCAGTTCTTCATTATACCGTAGATCTGAGACCTCTGTG  420
Sbjct  15138  ............................................................  15079

Query  421    CGTTCCATGGACTCAAGAGAGGCATTCTGTTTCCTGAAGACCAAAAGGAAGCCAAAAATC  480
Sbjct  15078  ............................................................  15019

Query  481    TTGGAAGAAGGACCTTTTCACAAAATTCACAAGGGTGTATACCAAGAGCTCCCTAACTCG  540
Sbjct  15018  ............................................................  14959

Query  541    GATCCTGCTGACCCAAACTCGGATCCTGCTGACCTAATTTTCCAAAAAGACTATCTTGAA  600
Sbjct  14958  ............................................................  14899

Query  601    TATGGTGAAATCCTGCCTTTTCCTGAGTCTCAGTTAGTAGAGTTTAAACAGTTCTCTACA  660
Sbjct  14898  ............................................................  14839

Query  661    AAACACTTCCAAGAATATGTAAAAAGGACAATTCCAGAATACGTCCCTGCATTTGCAAAC  720
Sbjct  14838  ............................................................  14779

Query  721    ACTGGAGGAGGCTATCTTTTTATTGGAGTGGATGATAAGAGTAGGGAAGTCCTGGGATGT  780
Sbjct  14778  ............................................................  14719

Query  781    GCAAAAGAAAATGTTGACCCTGACTCTTTGAGAAGGAAAATAGAACAAGCCATATACAAA  840
Sbjct  14718  ............................................................  14659

Query  841    CTACCTTGTGTTCATTTTTGCCAACCCCAACGCCCGATAACCTTCACACTCAAAATTGTG  900
Sbjct  14658  ............................................................  14599

Query  901    AATGTGTTAAAAAGGGGAGAGCTCTATGGCTATGCTTGCATGATCAGAGTAAATCCCTTC  960
Sbjct  14598  G...........................................................  14539

Query  961    TGCTGTGCAGTGTTCTCAGAAGCTCCCAATTCATGGATAGTGGAGGACAAGTACGTCTGC  1020
Sbjct  14538  ............................................................  14479

Query  1021   AGCCTGACAACCGAGAAATGGGTAGGCATGATGACAGACACAGATCCAG  1069
Sbjct  14478  .................................................  14430
```

#### Exon 5

Sequence ID: **Query_64009** Length: **27392** Number of Matches: **1**

**Range 1: 11934 to 12062** Graphics ▼ Next Match ▲ Previous

| Score | Expect | Identities | Gaps | Strand |
|---|---|---|---|---|
| 233 bits(258) | 1e-64 | 129/129(100%) | 0/129(0%) | Plus/Minus |

```
Query  1      ATCTTCTACAGTTGTCTGAAGATTTTGAATGTCAGCTGAGTCTATCTAGTGGGCCTCCCC  60
              ||||||||||||||||||||||||||||||||||||||||||||||||||||||||||||
Sbjct  12062  ATCTTCTACAGTTGTCTGAAGATTTTGAATGTCAGCTGAGTCTATCTAGTGGGCCTCCCC  12003

Query  61     TTAGCAGACCAGTGTACTCCAAGAAAGGCCTGGAACATAAAAAGGAACTCCAGCAACTTT  120
              ||||||||||||||||||||||||||||||||||||||||||||||||||||||||||||
Sbjct  12002  TTAGCAGACCAGTGTACTCCAAGAAAGGCCTGGAACATAAAAAGGAACTCCAGCAACTTT  11943

Query  121    TATTTTCAG  129
              |||||||||
Sbjct  11942  TATTTTCAG  11934
```

#### **Exon 6**

Sequence ID: **Query_48219** Length: **27392** Number of Matches: **1**

**Range 1: 5027 to 5750** Graphics ▼ Next Match ▲ Previous M

| Score | Expect | Identities | Gaps | Strand |
|---|---|---|---|---|
| 1293 bits(1433) | 0.0 | 721/724(99%) | 0/724(0%) | Plus/Minus |

```
Query  1     TCCCACCAGGATATTTGCGATATACTCCAGAGTCACTCTGGAGGGACCTGATCTCAGAGC  60
Sbjct  5750  ............................................................  5691

Query  61    ACAGAGGACTAGAGGAGTTAATAAATAAGCAAATGCAACCTTTCTTTCGGGGAATTTTGA  120
Sbjct  5690  ............................................................  5631

Query  121   TCTTCTCTAGAAGTTGGGCTGTGGACCTGAACTTGCAGGAGAAGCCAGGAGTCATCTGTG  180
Sbjct  5630  ............................................................  5571

Query  181   ATGCTCTGCTGATAGCACAGAACAGCACCCCCATTCTCTACACCATTCTCAGGGAGCAAG  240
Sbjct  5570  ..........................................................G.  5511

Query  241   ATGCAGAGGGCCAGGACTACTGCACTCGCACCGCCTTTACTTTGAAGCAGAAGCTAGTGA  300
Sbjct  5510  ...........................T...T............................  5451

Query  301   ACATGGGGGCTACACCGGGAAGGTGTGTGTCAGGGCCAAGGTCCTCTGCCTGAGTCCTG  360
Sbjct  5450  ............................................................  5391

Query  361   AGAGCAGCGCAGAGGCCTTGGAGGCTGCAGTGTCTCCGATGGATTACCCTGCGTCCTATA  420
Sbjct  5390  ............................................................  5331

Query  421   GCCTTGCAGGCACCCAGCACATGGAAGCCCTGCTGCAGTCCCTCGTGATTGTCTTACTCG  480
Sbjct  5330  ............................................................  5271

Query  481   GCTTCAGGTCTCTCTTGAGTGACCAGCTCGGCTGTGAGGTTTTAAATCTGCTCACAGCCC  540
Sbjct  5270  ............................................................  5211

Query  541   AGCAGTATGAGATATTCTCCAGAAGCCTCCGCAAGAACAGAGAGTTGTTTGTCCACGGCT  600
Sbjct  5210  ............................................................  5151

Query  601   TACCTGGCTCAGGGAAGACCATCATGGCCATGAAGATCATGGAGAAGATCAGGAATGTGT  660
Sbjct  5150  ............................................................  5091

Query  661   TTCACTGTGAGGCACACAGAATTCTCTACGTTTGTGAAAACCAGCCTCTGAGGAACTTTA  720
Sbjct  5090  ............................................................  5031

Query  721   TCAG  724
Sbjct  5030  ....  5027
```

#### **Exon 7**

**Sequence ID: Query_50973 Length: 27392 Number of Matches: 1**

**Range 1: 4047 to 4830** Graphics ▼ Next Match ▲ Previous I

| Score | Expect | Identities | Gaps | Strand |
|---|---|---|---|---|
| 1415 bits(1568) | 0.0 | 784/784(100%) | 0/784(0%) | Plus/Minus |

```
Query  1     TGATAGAAATATCTGCCGAGCAGAGACCCGGAAAACTTTCCTAAGAGAAAACTTTGAACA  60
Sbjct  4830  ............................................................  4771

Query  61    CATTCAACACATCGTCATTGACGAAGCTCAGAATTTCCGTACTGAAGATGGGGACTGGTA  120
Sbjct  4770  ............................................................  4711

Query  121   TGGGAAGGCAAAAAGCATCACTCGGAGAGCAAAGGGTGGCCCAGGAATTCTCTGGATCTT  180
Sbjct  4710  ............................................................  4651

Query  181   TCTGGATTACTTTCAGACCAGCCACTTGGATTGCAGTGGCCTCCCTCCTCTCTCAGACCA  240
Sbjct  4650  ............................................................  4591

Query  241   ATATCCAAGAGAAGAGCTCACCAGAATAGTTCGCAATGCAGATCCAATAGCCAAGTACTT  300
Sbjct  4590  ............................................................  4531

Query  301   ACAAAAAGAAATGCAAGTAATTAGAAGTAATCCTTCATTTAACATCCCCACTGGGTGCCT  360
Sbjct  4530  ............................................................  4471

Query  361   CGAGGTATTTCCTGAAGCCGAATGGTCCCAGGGTGTTCAGGGAACCTTACGAATTAAGAA  420
Sbjct  4470  ............................................................  4411

Query  421   ATACTTGACTGTGGAGCAAATAATGACCTGTGTGGCAGACACGTGCAGGCGCTTCTTTGA  480
Sbjct  4410  ............................................................  4351

Query  481   TAGGGGCTATTCTCCAAAGGATGTTGCTGTGCTTGTCAGCACCGCAAAAGAAGTGGAGCA  540
Sbjct  4350  ............................................................  4291

Query  541   CTATAAGTATGAGCTCTTGAAAGCAATGAGGAAGAAAAGGGTGGTGCAGCTCAGTGATGC  600
Sbjct  4290  ............................................................  4231

Query  601   ATGTGATATGTTGGGTGATCACATTGTGTTGGACAGTGTTCGGCGATTCTCAGGCCTGGA  660
Sbjct  4230  ............................................................  4171

Query  661   AAGGAGCATAGTGTTTGGGATCCATCCAAGGACAGCTGACCCAGCTATCTTACCCAATGT  720
Sbjct  4170  ............................................................  4111

Query  721   TCTGATCTGTCTGGCTTCCAGGGCAAAACAACACCTGTATATTTTTCCGTGGGGTGGCCA  780
Sbjct  4110  ............................................................  4051

Query  781   TTAG  784
Sbjct  4050  ....  4047
```

### SJEWS049193_D1

#### Exon 4

Sequence ID: **Query_31397** Length: **27392** Number of Matches: **2**

**Range 1: 14430 to 15498** Graphics ▼ Next Match ▲ Previou

| Score | Expect | Identities | Gaps | Strand |
|---|---|---|---|---|
| 1924 bits(2133) | 0.0 | 1068/1069(99%) | 0/1069(0%) | Plus/Minus |

```
Query  1      ATGGAGGCAAATCAGTGCCCCCTGGTTGTGGAACCATCTTACCCAGACCTGGTCATCAAT  60
Sbjct  15498  ............................................................  15439

Query  61     GTAGGAGAAGTGACTCTTGgagaagaaaacagaaaaaagctgcagaaaattcagagagac  120
Sbjct  15438  ............................................................  15379

Query  121    caagagaaggagagagTTATGCGGGCTGCATGTGCTTTATTAAACTCAGGAGGAGGAGTG  180
Sbjct  15378  ............................................................  15319

Query  181    ATTCGAATGGCCAAGAAGGTTGAGCATCCCGTGGAGATGGGACTGGATTTAGAACAGTCT  240
Sbjct  15318  ............................................................  15259

Query  241    TTGAGAGAGCTTATTCAGTCTTCAGATCTGCAGGCTTTCTTTGAGACCAAGCAACAAGGA  300
Sbjct  15258  ............................................................  15199

Query  301    AGGTGTTTTTACATTTTTGTTAAATCTTGGAGCAGTGGCCCTTTCCCTGAAGATCGCTCT  360
Sbjct  15198  ............................................................  15139

Query  361    GTCAAGCCCCGCCTTTGCAGCCTCAGTTCTTCATTATACCGTAGATCTGAGACCTCTGTG  420
Sbjct  15138  ............................................................  15079

Query  421    CGTTCCATGGACTCAAGAGAGGCATTCTGTTTCCTGAAGACCAAAAGGAAGCCAAAAATC  480
Sbjct  15078  ............................................................  15019

Query  481    TTGGAAGAAGGACCTTTTCACAAAATTCACAAGGGTGTATACCAAGAGCTCCCTAACTCG  540
Sbjct  15018  ............................................................  14959

Query  541    GATCCTGCTGACCCAAACTCGGATCCTGCTGACCTAATTTTCCAAAAAGACTATCTTGAA  600
Sbjct  14958  ............................................................  14899

Query  601    TATGGTGAAATCCTGCCTTTTCCTGAGTCTCAGTTAGTAGAGTTTAAACAGTTCTCTACA  660
Sbjct  14898  ............................................................  14839

Query  661    AAACACTTCCAAGAATATGTAAAAAGGACAATTCCAGAATACGTCCCTGCATTTGCAAAC  720
Sbjct  14838  ............................................................  14779

Query  721    ACTGGAGGAGGCTATCTTTTTATTGGAGTGGATGATAAGAGTAGGGAAGTCCTGGGATGT  780
Sbjct  14778  ............................................................  14719

Query  781    GCAAAAGAAAATGTTGACCCTGACTCTTTGAGAAGGAAAATAGAACAAGCCATATACAAA  840
Sbjct  14718  ............................................................  14659

Query  841    CTACCTTGTGTTCATTTTTGCCAACCCCAACGCCCGATAACCTTCACACTCAAAATTGTG  900
Sbjct  14658  ............................................................  14599

Query  901    AATGTGTTAAAAAGGGGAGAGCTCTATGGCTATGCTTGCATGATCAGAGTAAATCCCTTC  960
Sbjct  14598  G...........................................................  14539

Query  961    TGCTGTGCAGTGTTCTCAGAAGCTCCCAATTCATGGATAGTGGAGGACAAGTACGTCTGC  1020
Sbjct  14538  ............................................................  14479

Query  1021   AGCCTGACAACCGAGAAATGGGTAGGCATGATGACAGACACAGATCCAG  1069
Sbjct  14478  .................................................  14430
```

#### Exon 5

**Sequence ID: Query_4573  Length: 27392  Number of Matches: 1**

**Range 1: 11934 to 12062** Graphics  ▼ Next Match ▲ Previc

| Score | Expect | Identities | Gaps | Strand |
|---|---|---|---|---|
| 233 bits(258) | 1e-64 | 129/129(100%) | 0/129(0%) | Plus/Minus |

```
Query  1      ATCTTCTACAGTTGTCTGAAGATTTTGAATGTCAGCTGAGTCTATCTAGTGGGCCTCCCC  60
Sbjct  12062  ............................................................  12003

Query  61     TTAGCAGACCAGTGTACTCCAAGAAAGGCCTGGAACATAAAAAGGAACTCCAGCAACTTT  120
Sbjct  12002  ............................................................  11943

Query  121    TATTTTCAG  129
Sbjct  11942  .........  11934
```

### Exon 6

**Sequence ID: Query_55799  Length: 27392  Number of Matches: 1**

**Range 1: 5027 to 5750** Graphics  ▼ Next Match ▲ I

| Score | Expect | Identities | Gaps | Strand |
|---|---|---|---|---|
| 1293 bits(1433) | 0.0 | 721/724(99%) | 0/724(0%) | Plus/Minus |

```
Query  1     TCCCACCAGGATATTTGCGATATACTCCAGAGTCACTCTGGAGGGACCTGATCTCAGAGC  60
Sbjct  5750  ............................................................  5691

Query  61    ACAGAGGACTAGAGGAGTTAATAAATAAGCAAATGCAACCTTTCTTTCGGGGAATTTTGA  120
Sbjct  5690  ............................................................  5631

Query  121   TCTTCTCTAGAAGTTGGGCTGTGGACCTGAACTTGCAGGAGAAGCCAGGAGTCATCTGTG  180
Sbjct  5630  ............................................................  5571

Query  181   ATGCTCTGCTGATAGCACAGAACAGCACCCCCATTCTCTACACCATTCTCAGGGAGCAAG  240
Sbjct  5570  ..........................................................G.  5511

Query  241   ATGCAGAGGGCCAGGACTACTGCACTCGCACCGCCTTTACTTTGAAGCAGAAGCTAGTGA  300
Sbjct  5510  ...........................T...T............................  5451

Query  301   ACATGGGGGGCTACACCGGGAAGGTGTGTGTCAGGGCCAAGGTCCTCTGCCTGAGTCCTG  360
Sbjct  5450  ............................................................  5391

Query  361   AGAGCAGCGCAGAGGCCTTGGAGGCTGCAGTGTCTCCGATGGATTACCCTGCGTCCTATA  420
Sbjct  5390  ............................................................  5331

Query  421   GCCTTGCAGGCACCCAGCACATGGAAGCCCTGCTGCAGTCCCTCGTGATTGTCTTACTCG  480
Sbjct  5330  ............................................................  5271

Query  481   GCTTCAGGTCTCTCTTGAGTGACCAGCTCGGCTGTGAGGTTTTAAATCTGCTCACAGCCC  540
Sbjct  5270  ............................................................  5211

Query  541   AGCAGTATGAGATATTCTCCAGAAGCCTCCGCAAGAACAGAGAGTTGTTTGTCCACGGCT  600
Sbjct  5210  ............................................................  5151

Query  601   TACCTGGCTCAGGGAAGACCATCATGGCCATGAAGATCATGGAGAAGATCAGGAATGTGT  660
Sbjct  5150  ............................................................  5091

Query  661   TTCACTGTGAGGCACACAGAATTCTCTACGTTTGTGAAAACCAGCCTCTGAGGAACTTTA  720
Sbjct  5090  ............................................................  5031

Query  721   TCAG  724
Sbjct  5030  ....  5027
```

### Exon 7

Sequence ID: **Query_13815** Length: **27392** Number of Matches: **1**

**Range 1: 4047 to 4830** Graphics ▼ Next Match ▲ Previous

| Score | Expect | Identities | Gaps | Strand |
|---|---|---|---|---|
| 1415 bits(1568) | 0.0 | 784/784(100%) | 0/784(0%) | Plus/Minus |

```
Query  1     TGATAGAAATATCTGCCGAGCAGAGACCCGGAAAACTTTCCTAAGAGAAAACTTTGAACA  60
Sbjct  4830  ............................................................  4771

Query  61    CATTCAACACATCGTCATTGACGAAGCTCAGAATTTCCGTACTGAAGATGGGGACTGGTA  120
Sbjct  4770  ............................................................  4711

Query  121   TGGGAAGGCAAAAAGCATCACTCGGAGAGCAAAGGGTGGCCCAGGAATTCTCTGGATCTT  180
Sbjct  4710  ............................................................  4651

Query  181   TCTGGATTACTTTCAGACCAGCCACTTGGATTGCAGTGGCCTCCCTCCTCTCTCAGACCA  240
Sbjct  4650  ............................................................  4591

Query  241   ATATCCAAGAGAAGAGCTCACCAGAATAGTTCGCAATGCAGATCCAATAGCCAAGTACTT  300
Sbjct  4590  ............................................................  4531

Query  301   ACAAAAAGAAATGCAAGTAATTAGAAGTAATCCTTCATTTAACATCCCCACTGGGTGCCT  360
Sbjct  4530  ............................................................  4471

Query  361   CGAGGTATTTCCTGAAGCCGAATGGTCCCAGGGTGTTCAGGGAACCTTACGAATTAAGAA  420
Sbjct  4470  ............................................................  4411

Query  421   ATACTTGACTGTGGAGCAAATAATGACCTGTGTGGCAGACACGTGCAGGCGCTTCTTTGA  480
Sbjct  4410  ............................................................  4351

Query  481   TAGGGGCTATTCTCCAAAGGATGTTGCTGTGCTTGTCAGCACCGCAAAAGAAGTGGAGCA  540
Sbjct  4350  ............................................................  4291

Query  541   CTATAAGTATGAGCTCTTGAAAGCAATGAGGAAGAAAAGGGTGGTGCAGCTCAGTGATGC  600
Sbjct  4290  ............................................................  4231

Query  601   ATGTGATATGTTGGGTGATCACATTGTGTTGGACAGTGTTCGGCGATTCTCAGGCCTGGA  660
Sbjct  4230  ............................................................  4171

Query  661   AAGGAGCATAGTGTTTGGGATCCATCCAAGGACAGCTGACCCAGCTATCTTACCCAATGT  720
Sbjct  4170  ............................................................  4111

Query  721   TCTGATCTGTCTGGCTTCCAGGGCAAAACAACACCTGTATATTTTTCCGTGGGGTGGCCA  780
Sbjct  4110  ............................................................  4051

Query  781   TTAG  784
Sbjct  4050  ....  4047
```

**Description of Supplemental Tables**

Table S1A. Correlation between drug efficacy and SLFN11 expression in the GDSC.

Table S1B. SLFN11 expression levels in cell lines from the GDSC.

Table S1C. COSMIC ATM mutation status in the GDSC cell lines.

Table S1D. COSMIC BRCA1 mutation status in the GDSC cell lines.

Table S1E. COSMIC BRCA2 mutation status in the GDSC cell lines.

Table S1F. Mutational signatures calculated from pediatric patients with sarcoma.

Table S1G. Diagnoses for the patients in the IHC study.

Table S1H. IHC profiling of samples from pediatric patients with solid tumors.

Table S2A. Source, media, relative SLFN11 RNA and protein levels, and other information pertaining to the cell lines studied in this work.

Table S2B. ICH profiling of selected cell lines studied in this work.

Table S2C. EWS-FLI transcriptional targets.

Table S3. Preclinical *in vivo* testing of PARPi combinations.

Table S4A. Mean and median SLFN11 H-score and SLFN11 positivity as a function of disease progression.

Table S4B. Statistical analysis of SLFN11 H-score or SLFN11 positivity and patient outcomes.

Table S5A. List of 4960 genes that are differentially expressed in SLFN11 wild-type and knockout cell lines.

Table S6A. List of sensitive and resistance ES cell lines from the GDSC.

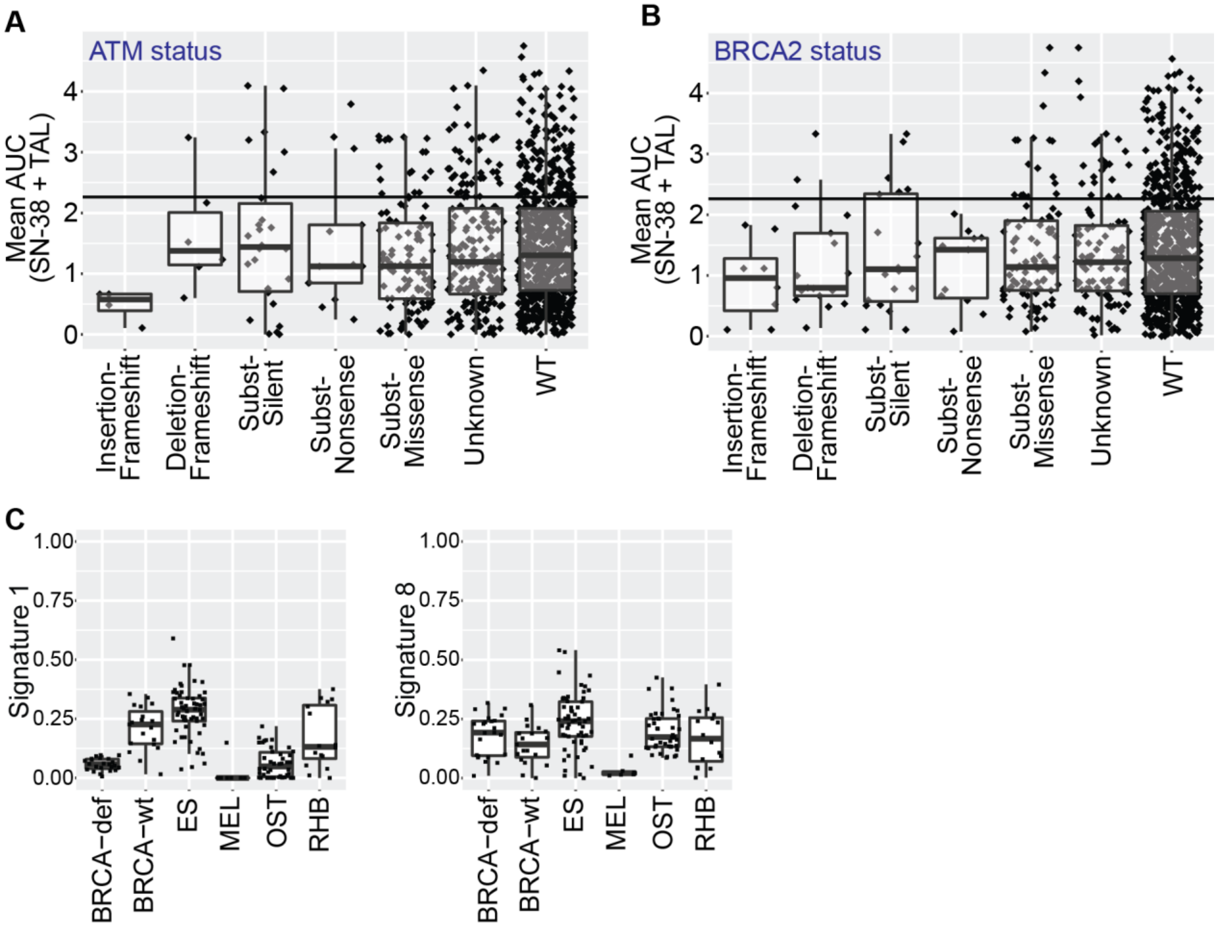


**Figure S1. (A, B)** Correlation between the mean AUC for SN-38 and TAL from the GDSC and ATM **(A)** and BRCA2 **(B)** mutational status as annotated in the COSMIC database. **(C)** Levels of mutational signatures 1 and 8 in samples from children with sarcoma. Melanoma, BRCA-wild-type, and BRCA-deficient samples are included for reference.

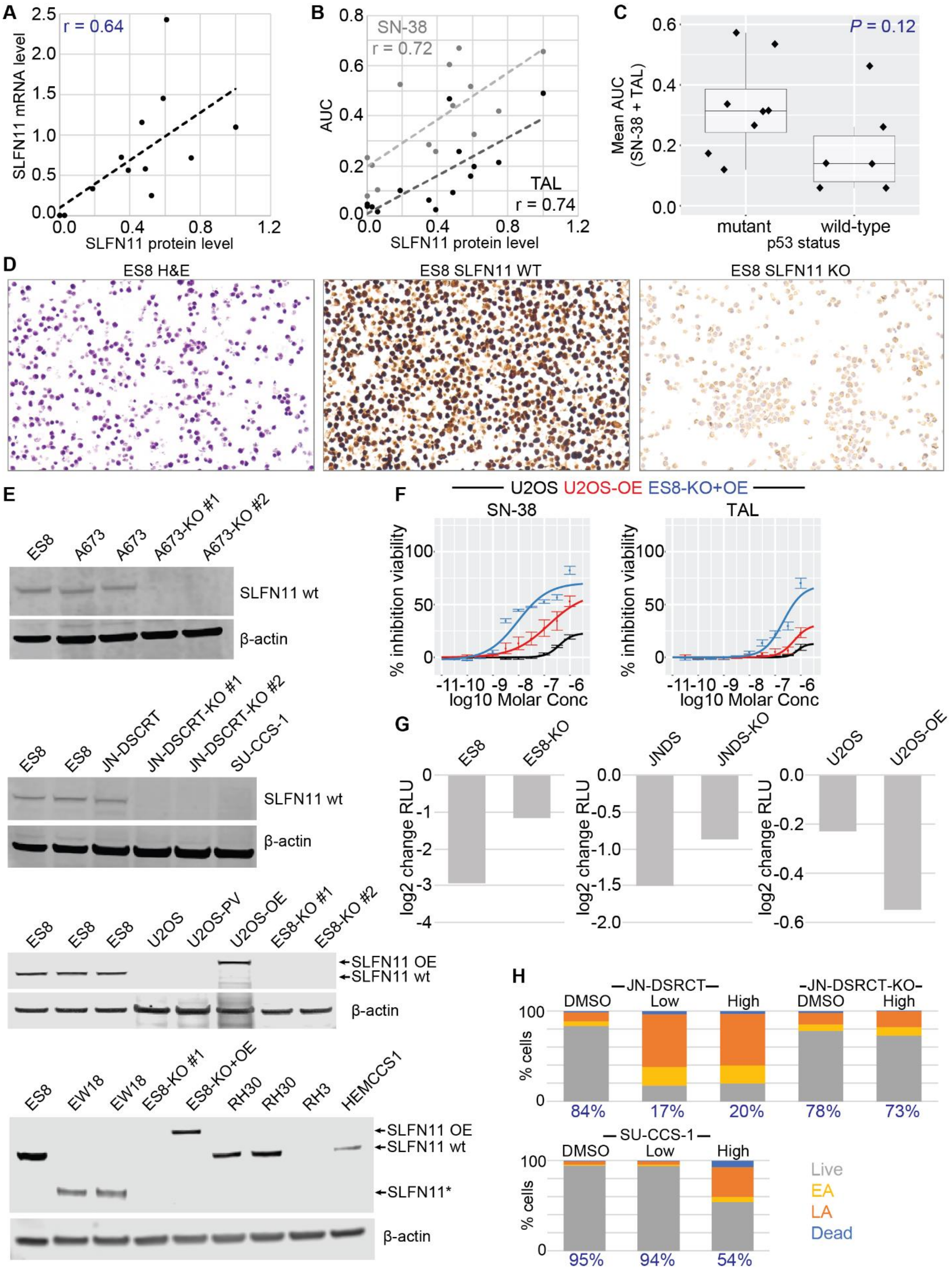
A
r = 0.64
SLFN11 mRNA level
SLFN11 protein level
B
SN-38
r = 0.72
TAL
r = 0.74
AUC
SLFN11 protein level
C
P = 0.12
Mean AUC (SN-38 + TAL)
mutant
wild-type
p53 status
D
ES8 H&E
ES8 SLFN11 WT
ES8 SLFN11 KO
E
ES8 A673 A673 A673-KO #1 A673-KO #2
SLFN11 wt
β-actin
ES8 ES8 JN-DSCRT JN-DSCRT-KO #1 JN-DSCRT-KO #2 SU-CCS-1
SLFN11 wt
β-actin
ES8 ES8 ES8 U2OS U2OS-PV U2OS-OE ES8-KO #1 ES8-KO #2
SLFN11 OE
SLFN11 wt
β-actin
ES8 EW18 EW18 ES8-KO #1 ES8-KO+OE RH30 RH30 RH3 HEMCCS1
SLFN11 OE
SLFN11 wt
SLFN11*
β-actin
F
U2OS U2OS-OE ES8-KO+OE
SN-38
TAL
% inhibition viability
log10 Molar Conc
G
ES8 ES8-KO
JNDS JNDS-KO
U2OS U2OS-OE
log2 change RLU
H
JN-DSRCT
JN-DSRCT-KO
DMSO Low High DMSO High
% cells
84% 17% 20% 78% 73%
SU-CCS-1
DMSO Low High
95% 94% 54%
Live
EA
LA
Dead

**Figure S2. (A, B)** Correlation between SLFN11 protein and mRNA levels **(A)** and the AUC for SN-38 and TAL **(B)** in our panel of 14 sarcoma cell lines. **(C)** Relationship between p53 status and the mean of the AUC for SN-38 and TAL in our panel of 14 sarcoma cell lines. **(D)** Validation of the SLFN11 IHC protocol in ES8 and ES8-KO cell lines. **(E)** Western blot confirmation of SLFN11 knock-out in ES8, JN-DSCRT, and A673 cells; and of SLFN11 over-expression in ES8 and U2OS cells. Other sarcoma cell lines are included for reference. **(F)** Dose-response curves (based on CTG assays) for SN-38 and TAL (with 72-h exposure) in U2OS (black), U2OS-OE (red), and ES8-KO+OE (blue) cells. n ≥ 2. **(G)** Change in $log_2$ CTG RLU at 72 h after exposure to 4 Gy radiation for knockout and over-expression models. n ≥ 2. **(H)** Flow cytometry assessment of cytotoxicity in JN-DSRCT, JN-DSCRT-KO, and SU-CCS-1 cells after 24 h exposure to “Low” (10 nM SN-38 + 10 nM TAL) and “High” (1 µM SN-38 + 1 µM TAL) drug combinations. The percentage of live cells is shown in blue. EA = early apoptosis. LA = late apoptosis. n ≥ 2.

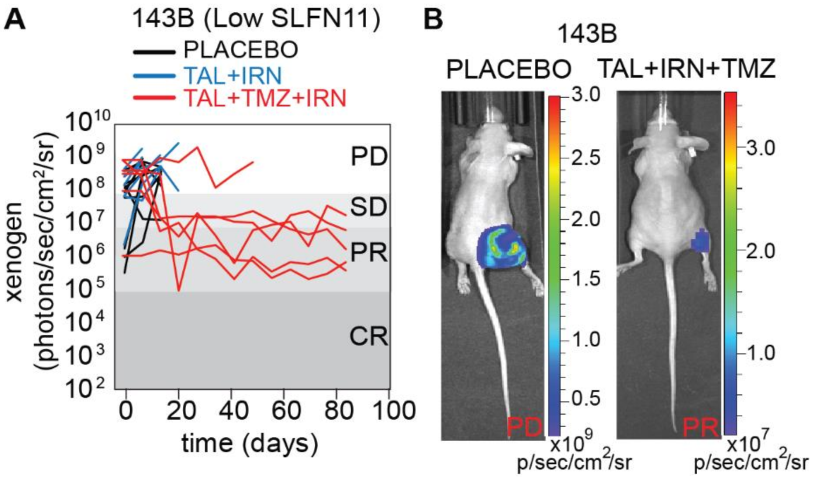


**Figure S3. (A)** Line plot of tumor burden over time as measured by bioluminescence for OST 143B xenografts. **(B)** Representative images from the study in **(A)**, showing progressive disease in the control group and partial response in the TAL+TMZ+IRN groups. PD, progressive disease; SD, stable disease; PR, partial response; CR, complete response.

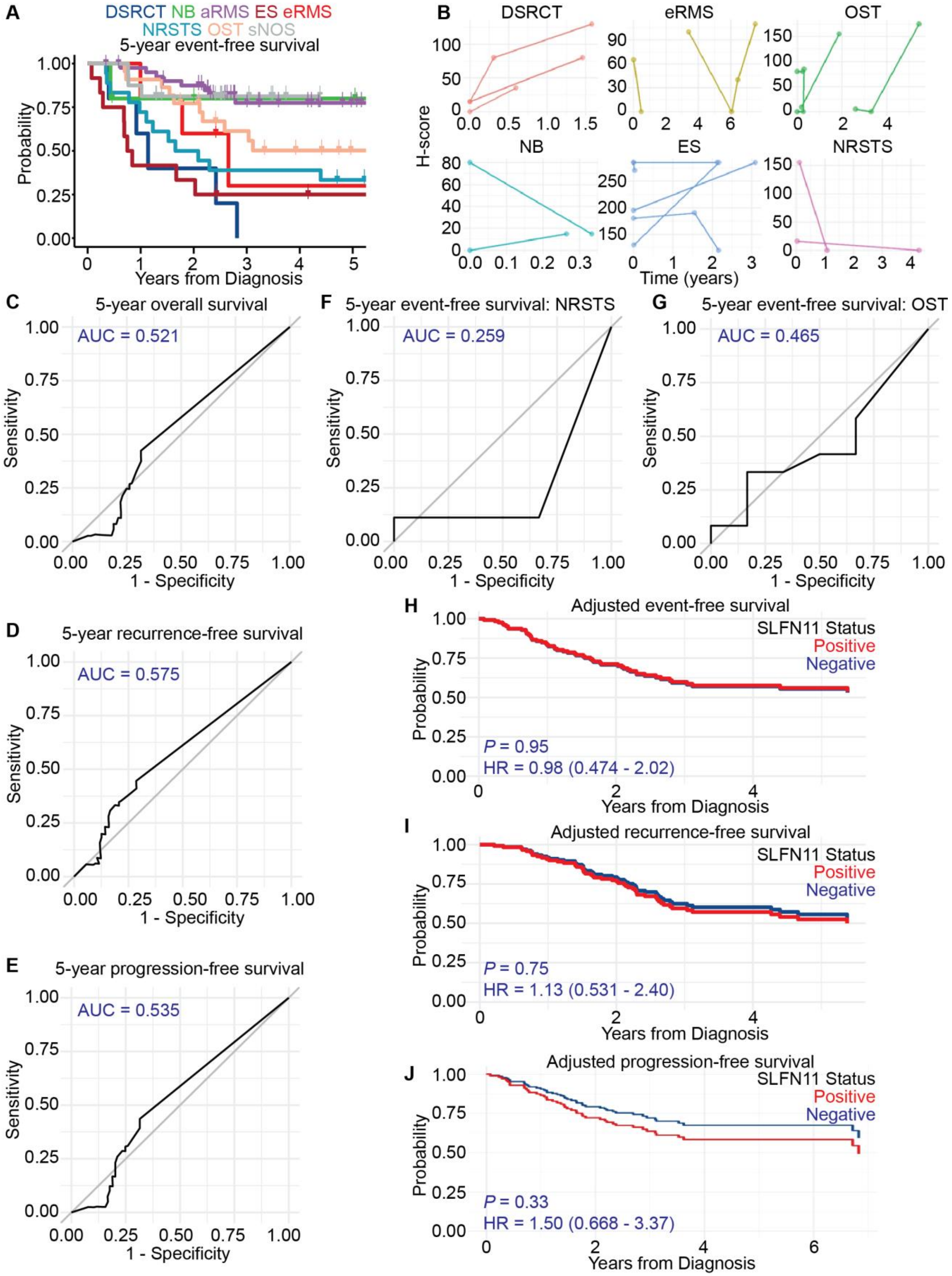
A
DSRCT NB aRMS ES eRMS
NRSTS OST sNOS
5-year event-free survival
Probability
Years from Diagnosis
B
DSRCT
eRMS
OST
NB
ES
NRSTS
H-score
Time (years)
C
5-year overall survival
AUC = 0.521
Sensitivity
1 - Specificity
F
5-year event-free survival: NRSTS
AUC = 0.259
G
5-year event-free survival: OST
AUC = 0.465
D
5-year recurrence-free survival
AUC = 0.575
E
5-year progression-free survival
AUC = 0.535
H
Adjusted event-free survival
SLFN11 Status
Positive
Negative
P = 0.95
HR = 0.98 (0.474 - 2.02)
I
Adjusted recurrence-free survival
P = 0.75
HR = 1.13 (0.531 - 2.40)
J
Adjusted progression-free survival
P = 0.33
HR = 1.50 (0.668 - 3.37)

**Figure S4. (A)** Five-year event-free survival rates by diagnosis for the patients with sarcoma in our IHC study. **(B)** SLFN11 H-score trajectories for 18 patients with at least 2 SLFN11 IHC measurements spanning different points in treatment, grouped by diagnosis. **(C–E)** ROC curves for 5-year overall, recurrence-free, and progression-free survival of all patients with sarcoma as a function of H-score. **(F–G)** ROC curves for 5-year event-free survival of NRSTS and OST patients, respectively, as a function of H-score. **(H–J)** Adjusted event-free, recurrence-free, and progression-free survival as a function of SLFN11 status after controlling for age, metastatic status, and disease.

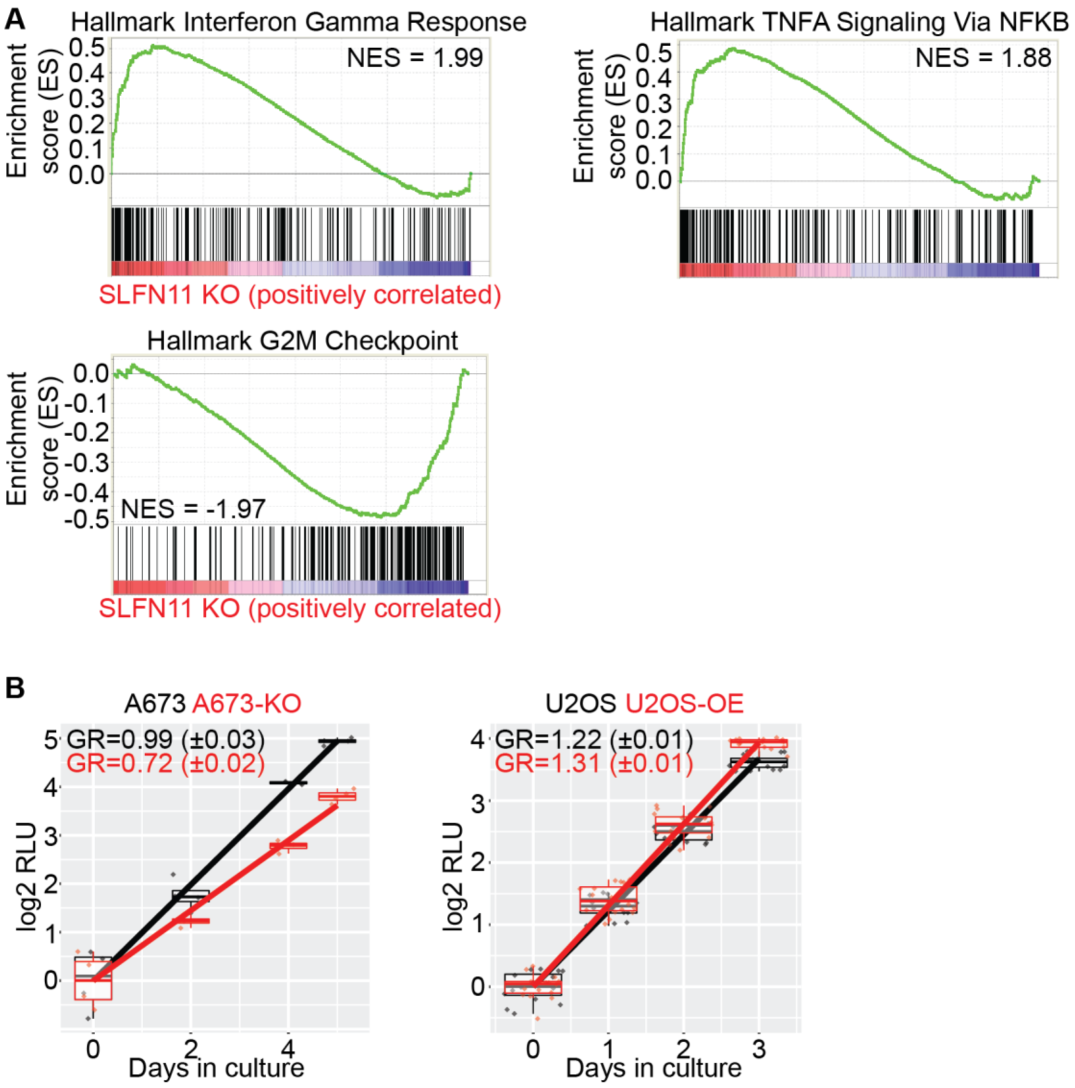


**Figure S5. (A)** Additional top-scoring GSEA Hallmark gene sets enriched in KO (top) and wild-type (bottom) lines (FDR q-value < 0.001). **(B)** Growth rate determination (based on CTG assays) in A673, A673-KO, U2OS, and U2OS-OE cells. GR = doublings/day. SEM reported in parentheses.

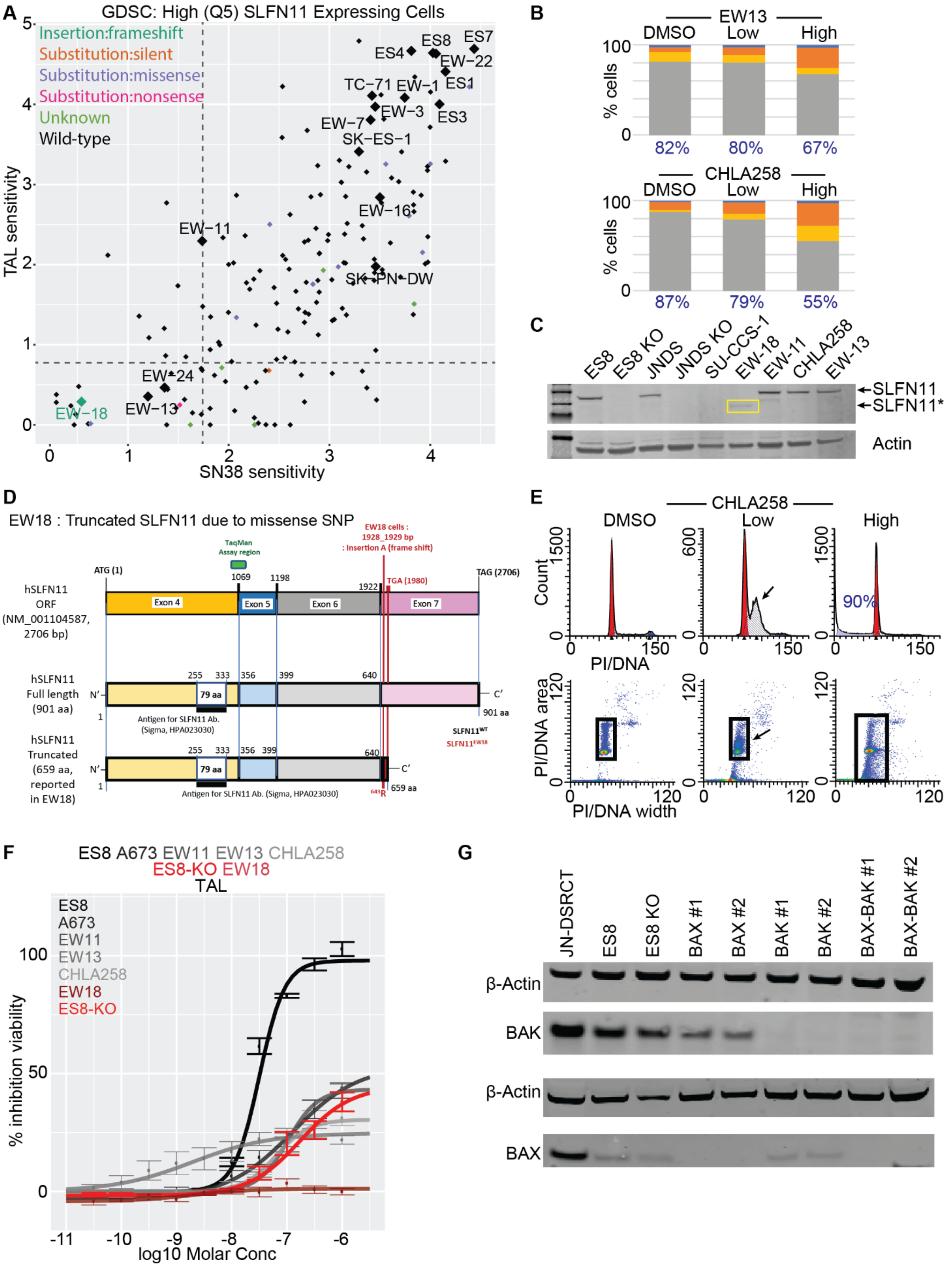
A
GDSC: High (Q5) SLFN11 Expressing Cells
Insertion:frameshift
Substitution:silent
Substitution:missense
Substitution:nonsense
Unknown
Wild-type
TAL sensitivity
SN38 sensitivity
B
EW13
CHLA258
DMSO
Low
High
% cells
82%
80%
67%
87%
79%
55%
C
ES8
ES8 KO
JNDS
JNDS KO
SU-CCS-1
EW-18
EW-11
CHLA258
EW-13
SLFN11
SLFN11*
Actin
D
EW18 : Truncated SLFN11 due to missense SNP
hSLFN11 ORF (NM_001104587, 2706 bp)
hSLFN11 Full length (901 aa)
hSLFN11 Truncated (659 aa, reported in EW18)
Antigen for SLFN11 Ab. (Sigma, HPA023030)
E
CHLA258
DMSO
Low
High
Count
PI/DNA
PI/DNA area
PI/DNA width
90%
F
ES8 A673 EW11 EW13 CHLA258
ES8-KO EW18
TAL
% inhibition viability
log10 Molar Conc
G
JN-DSRCT
ES8
ES8 KO
BAX #1
BAX #2
BAK #1
BAK #2
BAX-BAK #1
BAX-BAK #2
β-Actin
BAK
BAX

**Figure S6.** **(A)** Scatterplot of SN-38 vs TAL sensitivity for High ('Q5') SLFN11 expressing cell lines from the GDSC. Dotted lines represent the median drug activity observed for all cell lines in the database. ES cells are represented by diamonds. Colors depict the SLFN11 mutational status as annotated in the COSMIC database. **(B)** Flow cytometry assessment of cytotoxicity in EW13 and CHLA258 cells after exposure for 24 h to "Low" (10 nM SN-38 + 10 nM TAL) and "High" (1 µM SN-38 + 1 µM TAL) drug combinations. The percentage of live cells is shown in blue. EA = early apoptosis. LA = late apoptosis. n ≥ 1. **(C)** Western blot analysis of SLFN11 in resistant ES cell lines. ES8 and JN-DSCRT wild-type and knockout models, and SU-CCS-1 (SLFN11 null) are included for reference. **(D)** Diagram showing the EW-18 SLFN11 truncation and the recognition sites for the anti-SLFN11 antibody and TaqMan probes used in this study. **(E)** Cell cycle analysis of CHLA258 after exposure for 24 h to "Low" and "High" SN-38 + TAL. Arrows indicate the build-up of S-phase cells induced by the drug combination. **(F)** Dose-response curve (based on CTG assays) for TAL (with 72-h exposure) in ES8 cells (black), SLFN11-expressing resistant ES cell lines (grays), ES8 KO cells (red), and EW-18 cells (dark red). n ≥ 2. **(G)** Western blot confirming BAX and BAK KO expression in ES8 cells. JN-DSRCT was included as a positive control because it expressed high levels of both BAK and BAX.
